# Urea is a dynamic and functional component of mammalian breast milk

**DOI:** 10.64898/2026.09.28.755206

**Authors:** Liisa Veerus, Haipeng Sun, Axel A. Almet, Erica R. Levin, Rachana Rao Battaje, Vibeke S. Dam, Dareen Shah, Rachel Clerk, Brandon Chang, Melissa A. Woortman, Joyce E. Stuckey, Margot J. Shumaker, Yue Sandra Yin, Adrija Kundu, Meliza Talaue-Weeast, Zhan Gao, Valeria Cardenas, Xue-Song Zhang, Donald D. Nyangahu, Heather B. Jaspan, Michael Manhart, Qing Nie, Emily Barrett, Lene N. Nejsum, Maria Gloria Dominguez-Bello, Martin J. Blaser, the New Jersey Kids Study team

## Abstract

Urea is a constituent of human milk, yet its regulation and interactions with the infant gut microbiome have not been established. Combining three human cohorts, a mouse model, and microbiome analyses, we show that milk urea increases across lactation, is conserved across mammals, and is not explained by maternal serum urea or diet. Instead, we identify lactation-dependent remodeling of mammary urea metabolism and transport, including dynamic epithelial AQP3 localization consistent with regulated urea transfer into milk. Alongside urea, maternal milk and stool provide reservoirs of urea-metabolizing bacteria that help establish the infant “ureabiome”, with encoded urea-utilization capacity. This maternal–infant axis is context-dependent: infant formulas provide higher urea levels than human milk, while functional assays show that urea supports both commensal and opportunistic bacteria in a concentration-dependent manner. Together, these findings define a maternal nitrogen axis linking milk urea to infant gut microbiome assembly and function.

## Introduction

Lactation, a conserved mammalian trait^1,2^ provides newborns with nutrients, bioactive molecules, and cells essential for early development^3^. Human breast milk contains macronutrients, including carbohydrates, lipids, and proteins, along with immune components—antibodies, lactoferrin, lysozyme, and maternal immune cells^3,4^. Milk also carries diverse small molecules, including urea^5^, a nitrogenous end-product of protein catabolism^6^. Although measured in clinical settings from other bodily fluids, including urine, blood, sweat, and saliva, its presence in milk has been largely ignored except in dairy species^7–10^. Thus, whether urea is consistently present in human milk or in tractable experimental model subjects, such as mice, has been underexplored. The conservation of urea in mammalian milk^1,2^ however raises the fundamental question: why would a metabolic waste product appear in milk at all?

Answering this first requires understanding how urea reaches milk. Whether it simply equilibrates with circulating urea levels or is selectively regulated at the mammary epithelium was unknown. Candidate routes include urea-permeable aquaglyceroporins^11,12^ and canonical urea transporters^13,14^. Determining whether these pathways are regulated during lactation is therefore key to distinguishing controlled maternal transport from passive transfer.

If milk urea is physiologically regulated, its presence may serve a function beyond nitrogen excretion. Many milk constituents have been subject to strong selection to support infant development^1^. For example, human milk oligosaccharides (HMOs) are indigestible by infants yet nourish specific gut microbes as fermentable carbon sources^15,16^, shaping early-life microbial composition^17^. Considering that urea comprises a major share of the non-protein nitrogen in human milk^3^, it may play a parallel role in microbiome assembly by serving as a nitrogen source for urea-metabolizing gut bacteria^18^, including taxa linked to newborn health^19,20^, such as *Bifidobacterium longum* subspecies *infantis*^18^. Microbes that can utilize urea may thus have a competitive advantage within the early-life microbiome^21–23^, a period during which microbial states shape long-term developmental outcomes^24–26^. Commercially available infant formulas generally lack microbially targeted metabolites and do not report urea content, raising the possibility that formula feeding alters not only which microbes colonize the infant gut, but also the nutrient landscape that selects among them. Consistent with this idea, *B. infantis*, a key early-life symbiont, is enriched in breastfed infants, yet commonly diminished or absent in those fed formula^22^.

To shed light on these questions, we investigated the dynamics, regulation, and microbial consequences of milk urea. We first characterized its levels across lactation and examined mammary pathways that could regulate its transfer into milk. We then tested whether maternal stool and milk provide reservoirs of candidate urea-metabolizing bacteria for the infant gut and used metagenomics to define the functional capacity of the milk and infant “ureabiomes”. Finally, we compared physiological urea concentrations in human milk with those in commercial infant formulas and tested how increasing urea availability alters the growth and urease activity of infant-associated bacteria. Together, these analyses establish milk urea as a maternal metabolite with the capacity to modulate the early-life microbiome.

## Results

### Urea is a conserved, temporally regulated component of mammalian milk

We first addressed the presence and variation of urea concentrations in human milk by quantifying its temporal dynamics in a North American cohort (USA-1)^27^ across infant age, sequential months, and a single day. Breast milk urea concentrations differed significantly across four infant age groups (one-way ANOVA: F_3,34_ = 13.20, *P* = 7.08 × 10^-6^; **Fig. 1**a), with higher concentrations at 5–7 and 8–12 months than at 1 week (Tukey-adjusted *P* = 1.48 × 10^-5^ and 2.62 × 10^-4^, respectively), while the increase at 1–4 months relative to 1 week was borderline (*P* = 0.051). These age groups correspond to key lactation stages: early postpartum (1 week); established mature lactation with high milk intake (1–4 months); the transition to complementary feeding (5–7 months); and late lactation (8–12 months)^3^. Segmented linear regression identified a breakpoint at 20 weeks, with milk urea increasing through the first 5 months of life (slope = 0.231 mg/dL/week, *P* = 2.05 × 10⁻^5^) but showing no detectable change thereafter (slope = −0.047 mg/dL/week, *P* = 0.39; **Fig. 1**b). Based on this pattern, the two older infant groups were combined into a single 5–12-month category for all subsequent analyses. In sequential sampling performed one month apart, within-mother urea levels increased among mothers nursing 1–4-month-old infants (one-sample t-test: t = 2.80, df = 10, *P* = 0.019; **Fig. 1**c) but tended to decrease among mothers of 5–12-month-old infants (t = –1.93, df = 10, *P* = 0.082). These changes differed significantly between age groups (independent two-sample t-test: t = 3.37, df = 20, *P* = 0.003), indicating divergent month-to-month trajectories in milk urea across infancy. Within-day profiles also showed an infant age-dependent pattern (**Fig. 1**d): in the 1–4-month group, mean milk urea across 12:00, 18:00, and 24:00 was significantly higher than the 06:00 baseline (one-sample t-test: t = 2.60, df = 14, *P* = 0.021). No detectable within-day differences were observed for the 1-week and 5–12-month groups.

**Figure 1.**
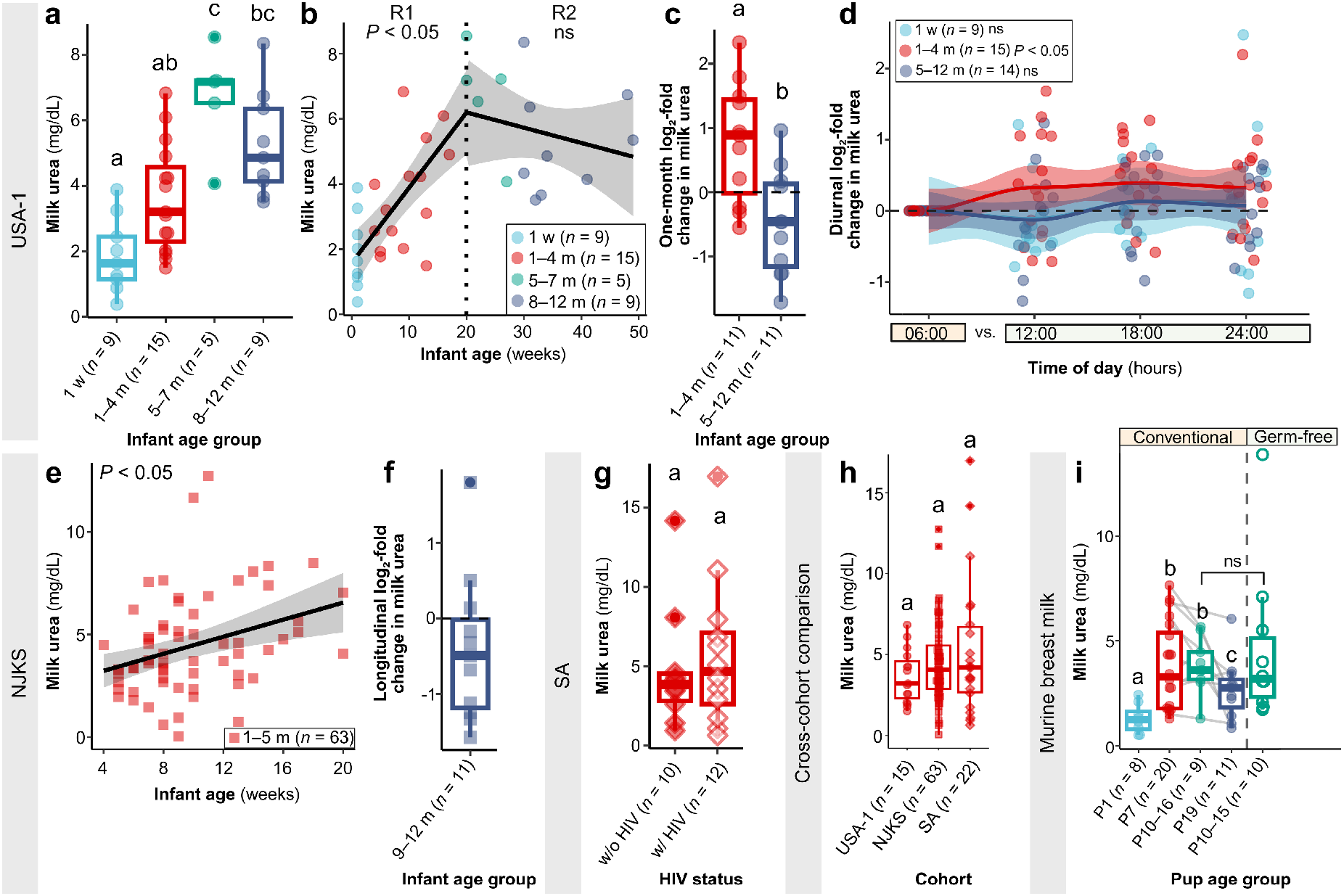
Milk urea is temporally regulated and conserved across human populations and mice. Each point represents a milk sample. Groups not sharing a letter differ significantly (*P* < 0.05 after multiple-comparison adjustment where applicable). (a) Milk urea levels across four infant age groups in 38 USA-1 mothers. (b) Segmented regression of milk urea levels versus infant age. Dashed line indicates the 20-week breakpoint and shading the 95% confidence interval. (c) One-month log_2_-fold change in milk urea levels in paired USA-1 samples. (d) Within-day log_2_-fold change in milk urea levels relative to 06:00 in USA-1 mothers sampled at 6-h intervals. (e) Association between infant age and milk urea levels in 63 NJKS mothers. (f) log_2_-fold change in milk urea levels from 1–5 to 9–12 months in 11 NJKS mothers with prolonged nursing. (g) Milk urea levels in 22 South Africa mothers living without or with HIV. (h) Milk urea levels across age-matched USA-1, NJKS, and SA cohorts. (i) Milk urea levels across lactation in conventional and germ-free mice. Gray lines connect repeated samples from the same dam. USA-1 – North America cohort 1; NJKS – New Jersey Kids Study; SA – South Africa cohort; P – postnatal day; w – week; m – months; w/o – without; w/ – with; ns – not significant.

Since the first 20 weeks of life represented a key phase of milk urea dynamics, we focused validation cohort comparisons on this age range. In the New Jersey Kids Study (NJKS) population, which included 63 mother–infant dyads sampled between 4 and 20 weeks postpartum, milk urea increased by 0.208 mg/dL/week (HC3-robust linear regression: t = 3.39, df = 61, *P* = 0.001; **Fig. 1**e), closely approximating the 0.231 mg/dL/week increase observed in the USA-1 cohort before 20 weeks postpartum (**Fig. 1**b). Although the within-mother decline in milk urea from 1–4 to 9–12 months did not reach statistical significance in NJKS (one-sample t-test: t = −1.38, df = 10, *P* = 0.197; **Fig. 1**f), urea levels decreased in 8 (72.7%) of 11 mothers. This directional pattern was consistent with USA-1 data, in which urea decreased in 7 (63.6%) of 11 mothers during later infancy (**Fig. 1**c). We also tested milk from mothers in South Africa (SA) nursing 4-week-old infants. Urea levels did not differ between mothers living with or without HIV (Wilcoxon rank-sum test: W = 52, *P* = 0.63; **Fig. 1**g), allowing us to consolidate the two groups. As a confirmatory analysis, we also repeated all within-cohort analyses after normalizing urea to total milk protein, with consistent results (**Extended Data Fig. 1**a–h). Finally, we compared age-matched samples across the three cohorts and found no detectable differences in milk urea (Kruskal–Wallis test: χ^2^ = 1.60, df = 2, *P* = 0.45; **Fig. 1**h), indicating comparable infant age-dependent concentrations across geographically distinct populations.

To assess whether the developmental pattern of milk urea observed in humans was more broadly conserved, we measured urea levels across lactation in control, antibiotic-treated, and germ-free mice. Antibiotic treatment did not detectably affect milk urea across mouse P7–19 (type III Wald test: χ^2^ = 0.50, df = 2, *P* = 0.78), with no age-by-treatment interaction (χ^2^ = 0.34, df = 4, *P* = 0.99), despite confirmed antibiotic-specific effects on the maternal gut microbiota (**Extended Data Fig. 2**a,b). Consolidating all conventional dams into a single group revealed that milk urea increased from postpartum colostrum (P1) to early (P7)^28^ and peak lactation (P10– 16)^29^, before declining at weaning (P19) (type II Wald test: χ^2^ = 30.20, df = 3, *P* = 1.25 × 10⁻^6^; **Fig. 1**i), recapitulating the temporal pattern observed in humans (**Fig. 1**a). At peak lactation, milk urea also did not differ between germ-free and age-matched conventional dams (F_1,17_ = 0.02, *P* = 0.89), indicating that an intact maternal microbiome is not required for urea incorporation into milk.

Collectively, these findings establish urea as a conserved component of milk that is characterized by a reproducible temporal pattern across geographically and clinically distinct human populations and mice, consistent with host-regulated control.

### Milk urea is uncoupled from maternal protein intake and serum urea levels, suggesting mammary epithelial transport

To investigate the physiological determinants of milk urea levels, we examined 34 NJKS mothers with paired serum and milk samples collected up to 20 weeks postpartum. The number of different animal-protein foods consumed in the previous 24 h was not associated with either milk urea (Gamma regression: milk urea ratio = 0.92 per additional food category, *P* = 0.44) or with serum urea levels (serum urea ratio = 1.22, *P* = 0.39; **Fig. 2**a) after adjustment for infant age and plant-protein consumption. Likewise, milk urea level was not associated with paired maternal serum urea after adjustment for infant age (Gamma regression: milk urea ratio = 1.00 per 1.00 mg/dL serum urea, *P* = 0.79; **Fig. 2**b), and milk–serum differences did not differ between 4–8 and 9–20 weeks postpartum (Wilcoxon W = 110, *P* = 0.25; **Fig. 2**c). Serum urea was not significantly associated with milk urea levels in mice either after adjustment for pup age (milk urea ratio = 1.05 per 1.00 mg/dL serum urea, *P* = 0.09; **Extended Data Fig. 2**c). Together, these findings indicate that variation in milk urea cannot be simply explained by maternal diet or circulating urea levels, suggesting that mammary glands may regulate urea transfer from the circulation into milk.

**Figure 2.**
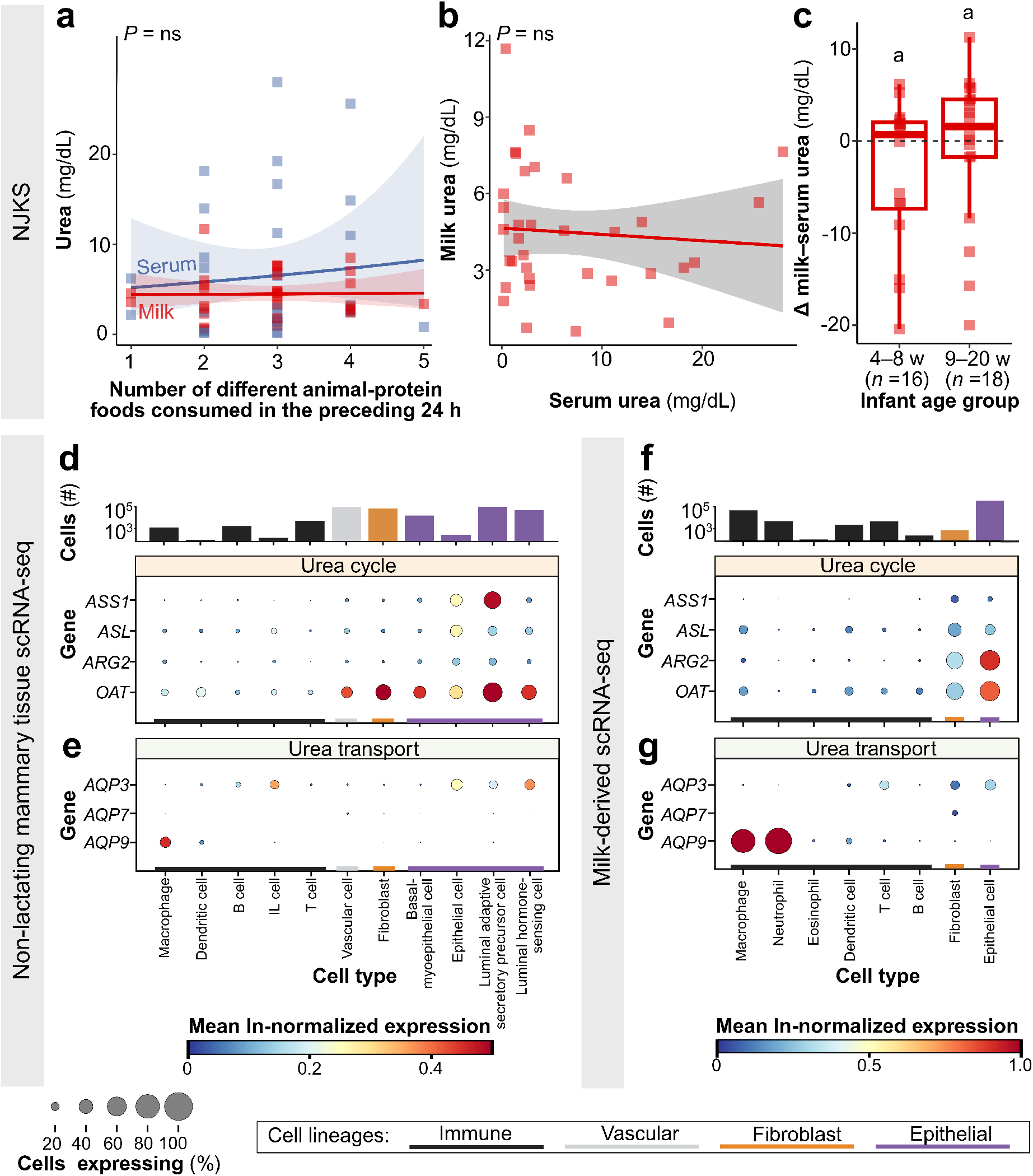
Human milk urea levels are not explained by maternal diet or serum urea, while urea-associated genes are partitioned across mammary cell types and enriched in milk-producing luminal epithelium. (a) Maternal serum and breast milk urea levels in 34 mothers according to the number of different animal-protein foods consumed within the previous 24 h. (b) Relationship between paired breast milk and maternal serum urea. (c) Paired breast milk–maternal serum urea difference by infant age group. Shared letters denote *P* > 0.05. Expression of (d,f) urea-cycle (*ASS1*, *ASL*) and associated ornithine-metabolism genes (*ARG2*, *OAT*), and (e,g) urea-permeable aquaglyceroporins (*AQP*) in (d,e) 36 non-lactating breast tissue samples from the Integrated Human Breast Cell Atlas^30^ and (f,g) cells derived from 59 human milk samples collected from 3 days to 90 weeks postpartum^31^. (d,f) Urea-metabolism genes are ordered according to pathway position, followed by associated ornithine-metabolism genes. (d–g) Cell types are grouped by lineage, with the number of cells per cell type shown above each dataset. Dot size indicates the percentage of cells expressing each gene, using a common scale across panels; color denotes mean natural-log-normalized expression, scaled independently within each dataset. NJKS – New Jersey Kids Study; w – weeks; ns – not significant.

We therefore asked whether the mammary gland is equipped for local urea production or for transepithelial transport of circulating urea into milk. Using publicly available single-cell RNA-sequencing datasets, we found that in healthy, non-lactating breast tissue^30^, the proximal urea-cycle genes *NAGS*, *CPS1*, and *OTC* were only minimally expressed (not shown), whereas downstream *ASS1*, *ASL*, *ARG2*, and the ornithine-metabolism gene *OAT* were more broadly detected (**Fig. 2**d). Among urea-permeable transporters, *AQP3* was substantially expressed in the luminal epithelial cells that line the alveolar space and participate in milk production (**Fig. 2**e). *AQP9* was concentrated in immune cells, while the canonical urea transporters *SLC14A1* and *SLC14A2* (encoding UT-B and UT-A, respectively) were scarcely or not at all detected (not shown).

Analysis of breast-milk-derived cells^31^ obtained during lactation revealed a similar pattern. Proximal urea-cycle genes remained sparsely expressed, whereas *ASS1*, *ASL*, *ARG2*, and *OAT* were detected, with particularly prominent *ARG2* and *OAT* expression in epithelial cells (**Fig. 2**f). *AQP3* was likewise expressed in milk-derived epithelial cells, whereas *AQP9* was predominantly restricted to immune cells, and *SLC14A1* and *SLC14A2* remained undetected (**Fig. 2**g).

This led us to focus specifically on *AQP3*. To assess the reproducibility of its epithelial enrichment, we examined seven independent scRNA-seq studies of non-lactating breast tissue^30,32–37^; luminal epithelial cells accounted for 69.2 ± 5.4% (mean ± SEM) of total *AQP3* expression. This epithelial cell enrichment was even more pronounced during lactation^31^, with 96.9 ± 0.9% of the total *AQP3* expression occurring in these cells.

In summary, these data provide evidence against a complete canonical urea cycle operating in the human breast. However, our findings support a role for transporter-mediated urea handling in mammary luminal epithelial cells, with *AQP3* emerging as the primary candidate.

### Lactation remodels murine mammary urea production and transport, and enriches epithelial AQP3

Because milk urea levels followed similar temporal dynamics in humans and mice (**Fig. 1**a,i), we used the mouse model, which permits direct sampling of lactating mammary tissue, to resolve mechanisms of urea handling that cannot be directly studied in humans. We performed bulk mammary RNA-seq, confirming the expected transcriptional transition into lactation, marked by strong stage-dependent regulation of *Lalba* and *B4galt1*, which together form the lactose synthase complex^38^, in both conventional and germ-free mice (**Fig. 3**a). Urea-cycle genes were also broadly regulated across reproductive stage (**Fig. 3**b). In conventional mice, all genes except *Cps1* showed significant stage-dependent regulation, whereas all genes were significant in germ-free mice (all adjusted *P* < 0.05). In both groups, *Ass1*, *Asl*, *Arg1*, and *Arg2* increased across lactation, while the pathway-initiating *Nags*–*Cps1*–*Otc* arm showed comparatively modest changes.

**Figure 3.**
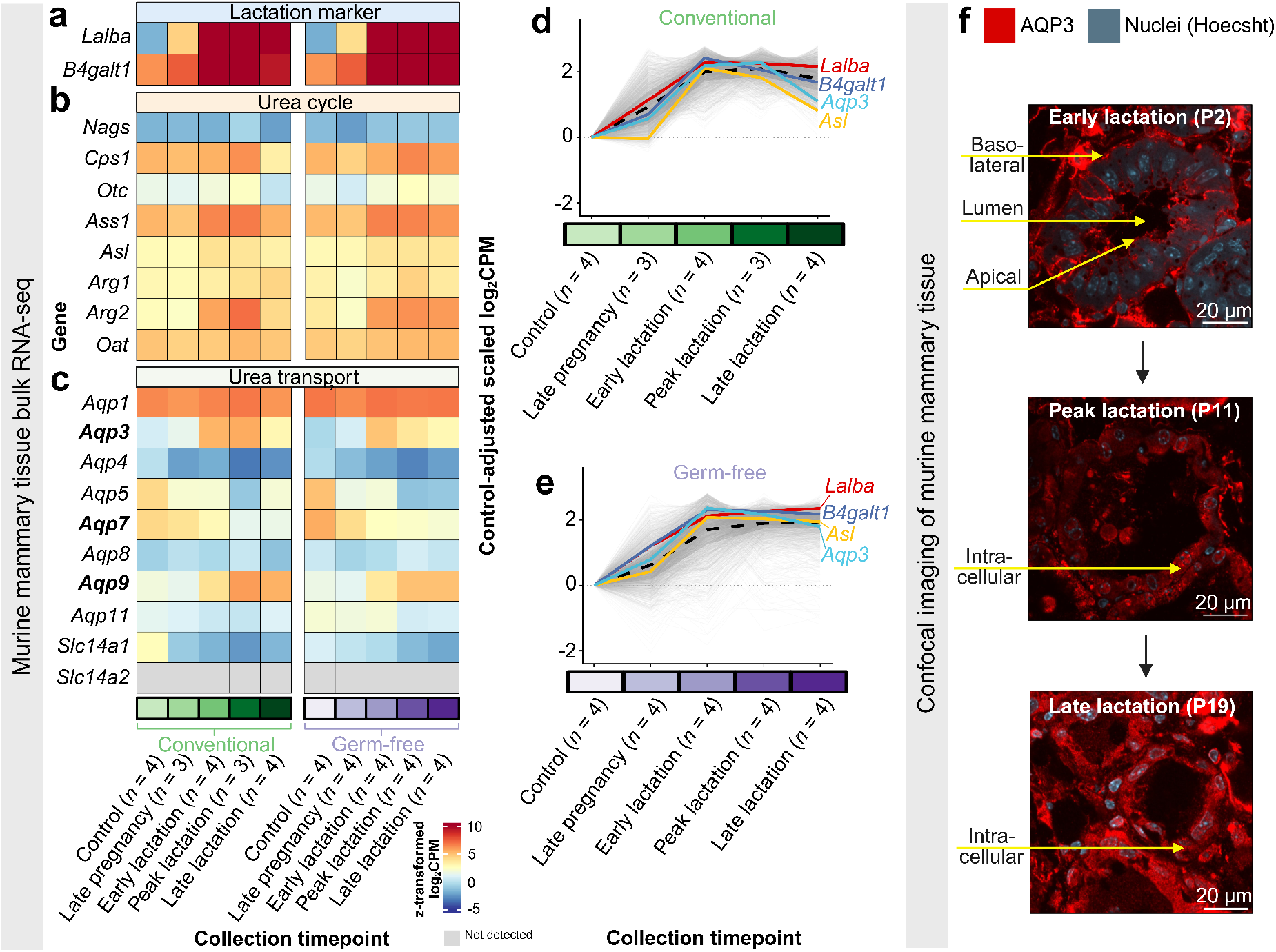
Lactation remodels murine mammary urea metabolism and transport, with dynamic AQP3 localization in the milk-producing mammary epithelium. (a– c) Heatmaps of fitted log_2_-transformed counts per million, standardized within each gene (z-transformed log_2_CPM), for genes encoding (a) lactation markers, (b) urea-cycle enzymes, and (c) aquaporins and canonical urea transporters in mammary tissue from conventional and germ-free female mice across non-pregnant/non-lactating control, late pregnancy, and early, peak, and late lactation. Genes encoding (c) urea-permeable aquaglyceroporins are in bold. (d,e) Expression trajectories of a 1,708-gene lactation-associated cluster showing significant reproductive stage-dependent regulation in both (d) conventional and (e) germ-free mice. Gray lines are individual genes; dashed line indicates the mean cluster trajectory. *Lalba* and *B4galt1*, canonical lactation genes^38^ that supported the annotation of the cluster as lactation-associated, are indicated together with the urea-cycle gene *Asl* and urea-permeable aquaglyceroporin *Aqp3*. (f) Representative confocal immunofluorescence images of murine mammary tissue during early, peak, and late lactation showing AQP3 localization (red) and nuclei (blue; Hoechst).

Urea transport machinery was similarly dynamic (**Fig. 3**c). All detected genes except *Aqp11* in conventional mice and *Aqp1* in germ-free mice were significantly regulated across reproductive stage (all adjusted *P* < 0.05). Expression of *Slc14a2* (encoding the canonical urea transporter UT-A) was not detected in any of the samples. Among aquaporins, we further focused on the urea-permeable aquaglyceroporins *Aqp3*, *Aqp7*, and *Aqp9*. Notably, *Aqp3* and *Aqp9* increased in expression during lactation, whereas *Aqp7* and the canonical urea transporter *Slc14a1* (UT-B) decreased. RT-qPCR confirmed the RNA-seq expression patterns of *Aqp3*, *Slc14a1*, and *Slc14a2* (**Extended Data Fig. 3**a–c), providing orthogonal validation of the bulk RNA-seq results.

We next performed transcriptome-wide clustering of mouse mammary tissue to identify broader reproductive-stage expression programs. Hierarchical clustering of 18,138 genes identified nine distinct temporal patterns, with one cluster comprising 1,708 genes increasing from late pregnancy through lactation before declining toward weaning, irrespective of microbiome status (**Fig. 3**d,e). This cluster contained *Lalba* (alpha-lactalbumin) and *B4galt1* (beta-1,4-galactosyltransferase 1), established components of the lactation program^38^. Among the urea-cycle genes, urea-permeable aquaporins, and canonical urea transporters examined (**Extended Data Fig. 3**d), *Asl* and *Aqp3* co-clustered with these canonical lactation genes, establishing their temporal regulation within the same lactation-associated transcriptional program.

Since bulk RNA-seq captures expression from all cell types in mammary tissue, we used the online Shiny app for the CRUK Cambridge Institute mouse mammary epithelial scRNA-seq atlas^39^ to resolve these findings within the mammary epithelium. The upstream *Nags*–*Cps1*–*Otc* arm was sparsely detected or absent, while downstream urea-cycle genes remained low in lactation-associated alveolar cells, providing further evidence against the activation of a complete urea cycle in the mammary epithelium. Among candidate transporters, *Aqp3* showed the clearest lactation-associated epithelial expression, while *Aqp9* and *Slc14a1* were sparse, and *Slc14a2* not detected. This closely mirrored the human scRNA-seq data (**Fig. 2**d–g).

Finally, we sought confirmation at the protein level by imaging the candidate urea transporters in lactating murine mammary tissue. AQP3 showed clear epithelial staining during lactation, including both basolateral and apical localization in early lactation, and increasing intracellular staining at later stages (**Fig. 3**f). By contrast, AQP9, UT-B, and UT-A1 showed no clear epithelial membrane localization during lactation (**Extended Data Fig. 3**e). Together, the human and murine findings identify AQP3, with its strong expression and lactation-dependency, as the strongest candidate mediator of regulated transepithelial urea transport into milk.

### Maternal microbiomes serve as reservoirs of urea-metabolizing bacteria for the infant gut

We next asked how candidate urea-metabolizing bacteria are distributed across maternal reservoirs and the infant gut. Mothers provide both a major reservoir of bacteria for the infant gut and, through breast milk, a source of urea that newly acquired bacteria may use as a nitrogen source. We therefore first characterized the maternal gut microbiome across pregnancy and early postpartum. Comparing 174 NJKS maternal stool samples, community composition varied modestly across reproductive stages (within-mother Bray–Curtis PERMANOVA: F= 0.94, R^2^ = 0.016, *P* = 0.0001; PERMDISP *P* = 0.015; **Fig. 4**a). The microbiome remained relatively stable across pregnancy, whereas the transition from third trimester to postpartum was the main significant pairwise shift (paired PERMANOVA: F_1,124_ = 0.92, R^2^ = 0.007, adjusted *P* = 0.001), consistent with peripartum restructuring of the maternal gut microbiota that coincides with the period of maternal–infant microbial transfer. In parallel, total bacterial load remained stable across pregnancy and postpartum (**Extended Data Fig. 4**a).

**Figure 4.**
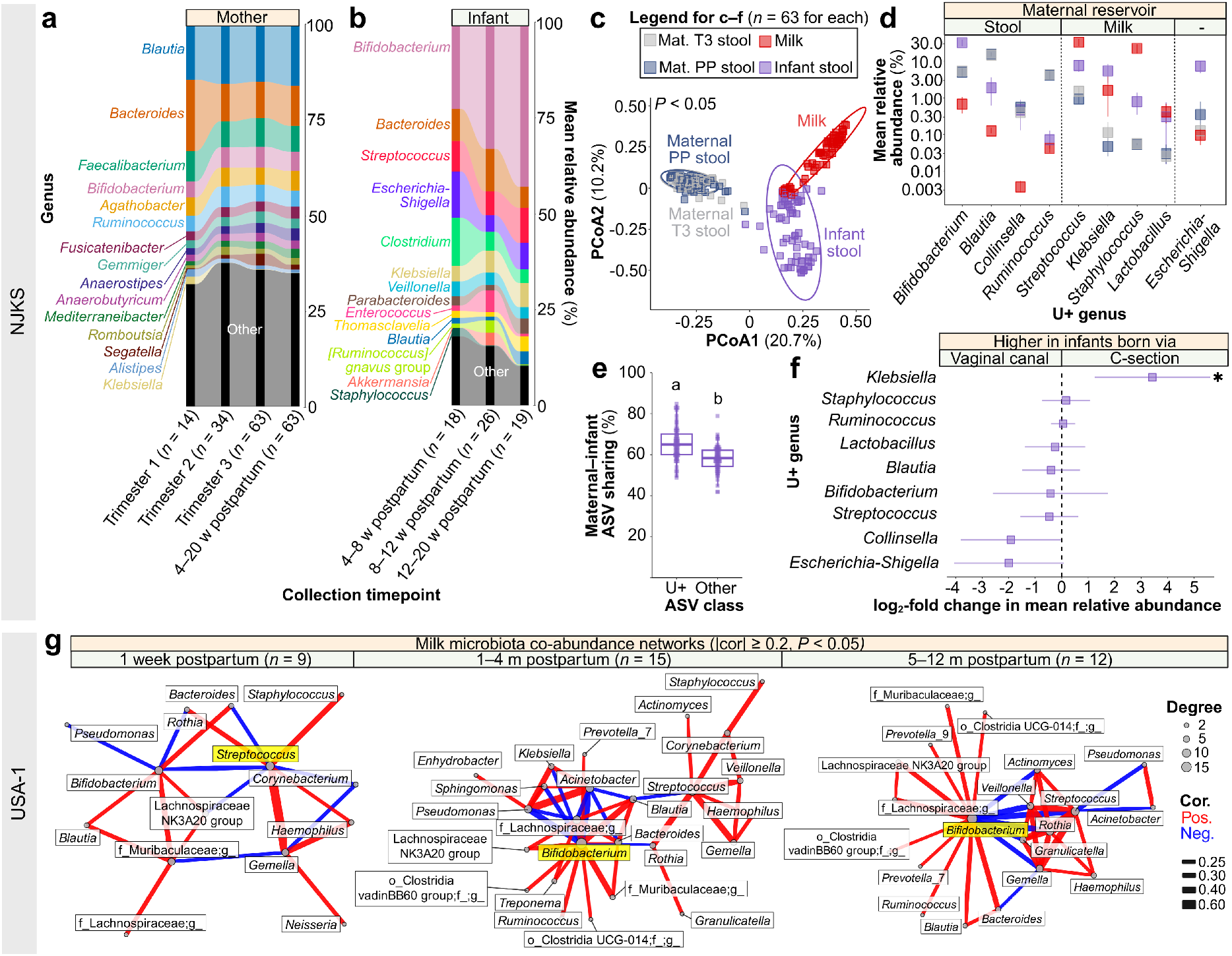
Maternal stool and milk are reservoirs of urea-metabolizing bacterial taxa for the infant gut. Genus-level composition of (a) the maternal stool microbiota across pregnancy and early postpartum, and (b) the infant stool microbiota across early postpartum. (a,b) The 10 most abundant genera overall or within any individual timepoint are indicated. (c–f) Analyses of 63 mother–infant dyads. (c) Bray–Curtis PCoA of maternal third-trimester and postpartum stool, milk, and infant stool at ASV level. Ellipses indicate 95% confidence regions. (d) Mean relative abundance of the nine candidate genera containing species or strains with reported or predicted urea-metabolizing capacity across maternal stool, milk, and infant stool, grouped by inferred maternal reservoir. Genera are ordered by decreasing mean relative abundance in infant stool within each reservoir group. The y-axis is log_10_-scaled; error bars indicate bootstrapped 95% confidence intervals. (e) Sharing of candidate urea-metabolizing ASVs and other genus-assigned ASVs between maternal third trimester and infant stool. Different letters indicate significant differences. (f) Association between birth mode and the relative abundance of the nine candidate genera in infant stool, adjusted for infant age, formula exposure, and milk urea levels. Error bars indicate 95% confidence interval; * indicates Benjamini–Hochberg-adjusted *P* < 0.05. (g) Milk microbiota co-abundance networks from 36 USA-1 mothers. The genus with the highest degree centrality in each postpartum window is highlighted in yellow. NJKS – New Jersey Kids Study; USA-1 – North America cohort 1; ASV – amplicon sequence variant; PCoA – principal coordinate analysis; T3 – third trimester; PP – postpartum; U+ – candidate urea-metabolizing genera; w – weeks; m – months.

Infant gut community composition remained broadly stable across 4–20 weeks postpartum (Bray–Curtis PERMANOVA: F_2,60_ = 1.39, R^2^ = 0.044, *P* = 0.091; PERMDISP *P* = 0.146; **Fig. 4**b), with a nominal difference between 4–8 and ≥ 12 weeks (F_1,37_ = 1.99, R^2^ = 0.054, *P* = 0.027, adjusted *P* = 0.080).

At the whole-community level, infant gut microbiota was distinct from the three principal maternal bacterial reservoirs—third-trimester and postpartum stool, and milk (ASV-level Bray–Curtis PERMANOVA: F_3,248_ = 31.27, R^2^ = 0.27, *P* = 0.0001; pairwise R^2^ = 0.22, 0.22, and 0.16, respectively; all adjusted *P* < 0.001; **Fig. 4**c). Total bacterial load was also significantly lower in infant than maternal stool, further distinguishing the infant gut from the maternal gut in both community composition and bacterial abundance (**Extended Data Fig. 4**b).

Next, we focused on nine genera containing species or strains with reported or predicted urea-metabolizing potential^18,40–43^ and asked how they were distributed across the principal maternal reservoirs—third-trimester and postpartum stool, and milk—and the infant gut (**Fig. 4**d). *Blautia*, *Ruminococcus*, *Bifidobacterium*, and *Collinsella* were enriched in maternal stool relative to milk, whereas *Staphylococcus*, *Streptococcus*, *Klebsiella*, and *Lactobacillus* were enriched in milk (all adjusted *P* < 0.01). *Escherichia*-*Shigella* did not differ detectably among maternal compartments. The population structure of the infant gut microbiome was reshaped compared to maternal communities, with pronounced expansion of *Bifidobacterium* and *Escherichia*-*Shigella*, depletion of *Blautia*, *Ruminococcus*, and *Collinsella* relative to maternal stool, and lower *Staphylococcus* and *Streptococcus* abundance relative to milk (all adjusted *P* < 0.001). Thus, the early-life ureabiome draws from distinct maternal reservoirs before substantial taxon-specific restructuring in the infant gut.

We then asked whether these nine taxa also showed evidence of maternal– infant sharing at finer resolution. Using third-trimester maternal stool as a proxy for gut bacteria available at delivery^44,45^, we found significantly greater maternal–infant ASV identity among nine candidate urea-metabolizing genera than among other taxa (65.6 ± 1.0% versus 58.1 ± 0.8%; paired Wilcoxon test, *P* = 3.45 × 10⁻^7^; **Fig. 4**e). We next asked whether this sharing was specific to biological mother–infant pairs. Urea-metabolizing ASV overlap was similarly high with unrelated mothers (64.9 ± 0.9%; own versus unrelated, *P* = 0.52), indicating that these candidate taxa are broadly conserved across mothers and consistently represented in the early-life ureabiome.

Birth mode further shaped these candidate genera in the infant gut (**Fig. 4**f): after accounting for infant age, formula exposure, and milk urea, *Klebsiella* was enriched in Cesarean-delivered infants (log_2_-fold change = 3.43, adjusted *P* = 0.028), whereas no other candidate genus differed by delivery mode (all adjusted *P* ≥ 0.19).

Because milk urea showed its strongest age-dependent separation in the longitudinal USA-1 cohort, we next investigated whether distinct milk “lactotypes” emerged across the same stages of lactation. Co-abundance networks revealed substantial reorganization of the milk microbiota across lactation (**Fig. 4**g). Although mean degree remained similar across age groups (3.87, 3.83, and 4.00), neighborhood connectivity increased (6.65, 7.70, and 10.34), indicating greater organization around highly connected taxa later in lactation. The dominant hubs also shifted with infant age: *Streptococcus* was the highest-degree genus at 1 week postpartum, whereas *Bifidobacterium* was the main hub at both 1–4 and 5–12 months, accompanied by a marked increase in betweenness centrality (0.03 to 0.58).

Our findings reveal a dynamic maternal–infant microbial axis in which a conserved set of candidate urea-metabolizing bacteria partitions across complementary maternal reservoirs before undergoing taxon-specific restructuring in the infant gut.

### Milk and infant gut microbiomes harbor distinct ureabiomes with conserved urea-utilization capacity

We then examined whether the milk and infant gut microbiomes contain the taxonomic and functional capacity to metabolize urea. In USA-1 milk metagenomes, taxa with urea-metabolizing potential comprised 10.8% of the microbial community on average and were dominated by *Staphylococcus*, *Streptococcus*, *Pseudomonas*, and *Acinetobacter* (**Fig. 5**a); in unrelated four-week-old SA infant stool, the corresponding ureabiome comprised 6.6% of the microbial community and was instead dominated by *Klebsiella*, *Streptococcus*, *Enterobacteriaceae*, *Bifidobacterium*, and *Escherichia* (**Fig. 5**b). Although derived from independent cohorts, these metagenomic profiles supported the NJKS observation that candidate urea-metabolizing taxa partition by niche (**Fig. 4**d), with some genera preferentially associated with milk and others with stool/gut environments. Notably, 6 (66.7%) of the 9 literature-defined candidate urea-metabolizing genera^18,40–43^ were independently represented across these metagenomic ureabiomes, further supporting their inclusion as candidate urea-metabolizing taxa.

**Figure 5.**
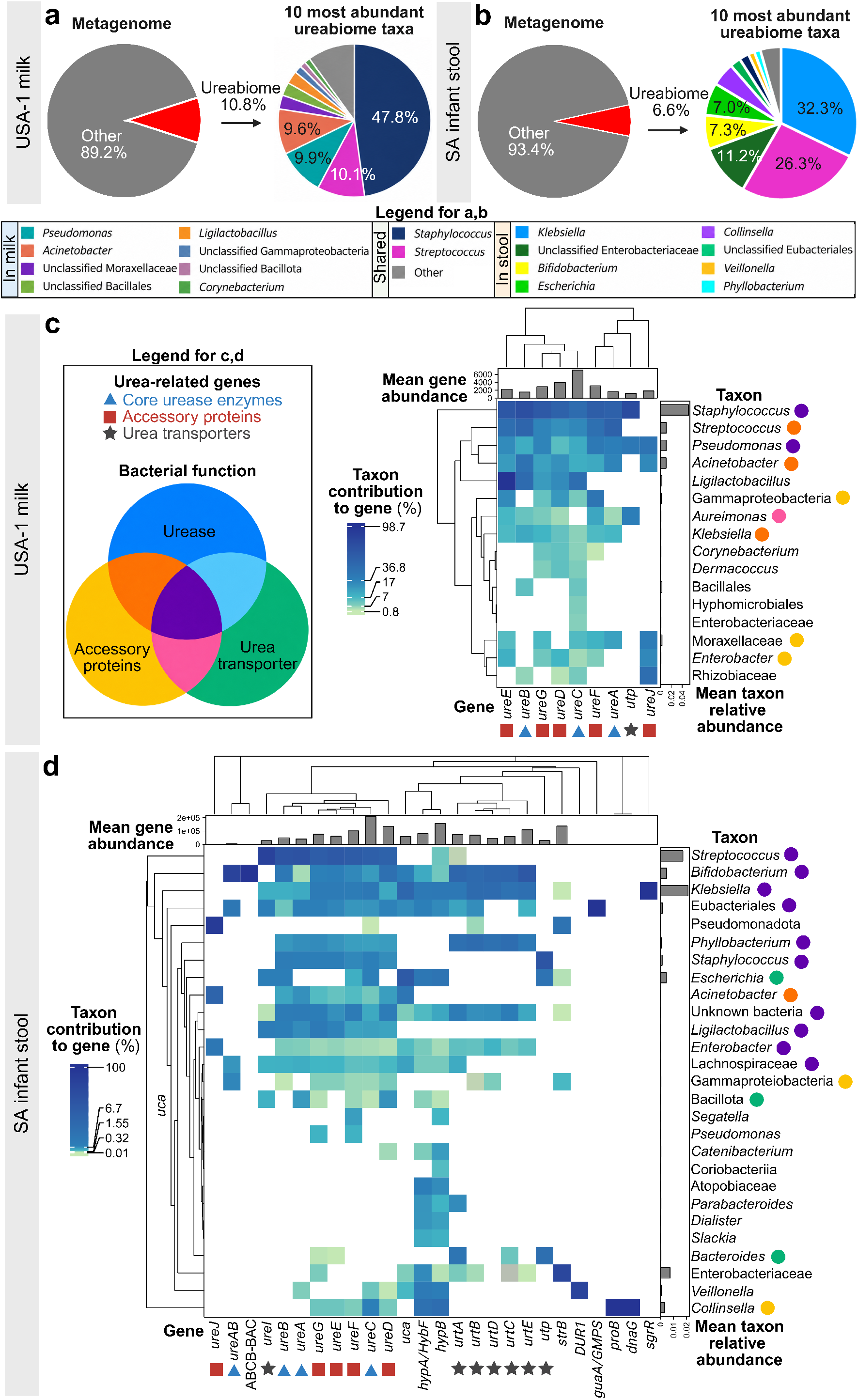
Taxonomy and functional capacity of the ureabiome in infant stool and human milk metagenomes. Proportion and taxonomic composition of the ureabiome in (a) USA-1 human milk metagenomes collected from 32 mothers across the first postpartum year and (b) infant stool metagenomes from 19 four-week-old SA infants. The 10 most abundant ureabiome taxa within each sample type are indicated and displayed in the legend in descending order of abundance. (c,d) Taxonomic contributions to urea-associated genes, based on KEGG Orthology annotations, in (c) USA-1 milk and (d) SA infant stool metagenomes. Taxa are shown at the genus level where available; otherwise, the highest resolved taxonomic rank is reported. Color intensity indicates the percentage contribution of each taxon to a given urea-associated gene; white indicates no detectable contribution. Bar plots above the heatmaps show the mean read abundance of each gene, whereas bar plots on the right show mean taxon relative abundance. Symbols below the heatmaps denote core urease structural genes (*ureA*–*ureC* and *ureAB*), urease accessory genes (*ureD*–*ureG* and *ureJ*), and urea transport genes, including the acid-activated urea channel *ureI*^46^, the ATP-dependent urea transporter *urtA*–*urtE*^47^, and the single-component urea transporter *utp*^48^. Symbols beside taxa denote inferred core urease, accessory, and urea transport capacity, assigned for complete core urease machinery, ≥ 80% of accessory genes, and/or at least one complete urea transport mechanism. USA-1 – North America cohort 1; SA – South Africa cohort.

We next resolved the taxonomic contributions to genes involved in urea metabolism. Both milk and infant stool metagenomes contained core urease structural genes, urease accessory machinery, and multiple urea transport systems, including the acid-activated channel *ureI*^46^, ATP-dependent *urtABCDE* transporters^47^, and the single-component transporter *utp*^48^ (**Fig. 5**c,d). These functions were distributed across multiple taxa rather than confined to a single lineage. In infant stool, *Streptococcus*, *Bifidobacterium*, and *Klebsiella* encoded the complete core urease machinery together with accessory and urea transport capacity (**Fig. 5**d). Combined with their high relative abundance, these features highlighted their potential for urea utilization in the early-life gut.

Because urease activity provides ammonia that can be incorporated into central nitrogen metabolism, we next focused specifically on infant stool metagenomes, evaluating urea-associated functions using Clusters of Orthologous Groups (COGs). We first confirmed the presence of urease structural and accessory genes and urea transporters across infants (**Extended Data Fig. 5**). Downstream nitrogen-utilizing functions consistent with urea hydrolysis and subsequent nitrogen assimilation, including allophanate hydrolases and branched-chain amino acid transport systems^49^, were variably detected across subjects.

These data establish that both milk and the early-life infant gut harbor ureabiomes that are taxonomically distinct, yet share the encoded capacity to access and metabolize urea.

### Infant feeding alters urea exposure, with potential consequences for early-life bacteria

Having established maternal sources of both urea and urea-metabolizing bacteria, we finally asked whether this system could be challenged by feeding mode. Urea concentrations differed markedly between commercial infant formulas and human milk across lactation (Kruskal–Wallis: χ^2^ = 87.71, df = 10, *P* = 1.5 × 10⁻^14^; **Fig. 6**a; **Extended Data Fig. 1**i), with five of eight formulas containing more urea than human milk at all three infant ages examined. We then asked whether these differences reflected the primary ingredients themselves or the processing methodology. Urea concentrations varied strongly among animal milks and plant-based beverages (**Fig. 6**b), with the highest levels in animal milks. Heating of milk (e.g., to temperatures relevant to warming before feeding or pasteurization) had little effect on urea in aqueous solution or in human milk and reduced it only after boiling cow milk (**Fig. 6**c). Given that many commercial infant formulas are cow-milk based, these findings indicate that their elevated urea concentrations reflect the urea-rich composition of the source milk.

**Figure 6.**
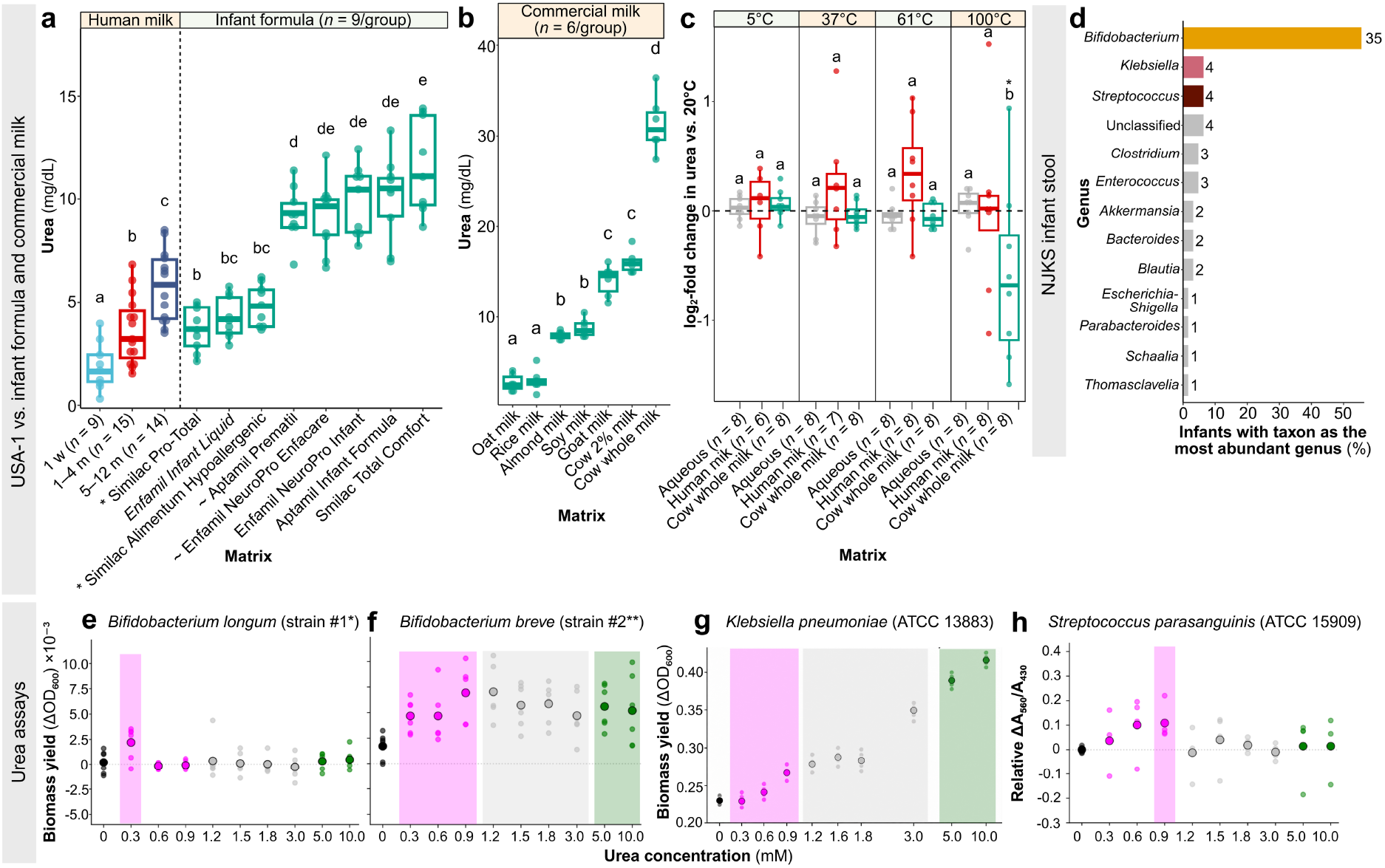
Urea availability differs across infant feeding sources and elicits concentration-specific functional responses in infant-associated bacteria. (a) Urea concentrations in human milk from the USA-1 cohort (Fig. 1a) grouped into three infant-age categories (left), and commercially available infant formulas (right). Formula displayed in italics is a liquid product; all others are powder-based. * denotes formulas in which the first ingredient after water is corn maltodextrin; the remaining formulations have non-fat milk as their primary ingredient. Products formulated specifically for premature infants indicated with ∼; all other formulas are intended for full-term infants. (b) Urea concentrations in commercial plant-based beverages and goat and cow milk. (a,b) Groups not sharing a letter differ significantly (all adjusted *P* < 0.05). (c) Temperature-dependent stability of urea in aqueous solution, human breast milk, and cow whole milk, shown as log_2_-fold change relative to the corresponding 20°C handling condition. Within each temperature, groups not sharing a letter differ significantly between matrices; * denotes a significant within-matrix change from the matched 20°C control (all adjusted *P* < 0.05). (d) Dominant genera in stools from 63 NJKS infants. Colored bars indicate the genera and the number of infants in which it was the most abundant genus *Bifidobacterium*, *Klebsiella* and *Streptococcus* are highlighted as genera represented in subsequent functional assays. (e–g) Biomass yield or (h) urease activity of candidate urea-metabolizing bacterial strains (**Extended Data Table 3**) across urea concentrations. Black denotes the no-urea control, pink physiological concentrations found in milk, grey infant-formula-range, and green supraphysiological levels comparable to cow milk. Shading indicates concentrations at which bacterial activity was significantly (*P* < 0.05) higher than the no-urea control. USA-1 – North America cohort 1; NJKS – New Jersey Kids Study; w – week; m – months.

We then returned to the NJKS infant microbiome to identify bacteria most likely to respond to these differences. Remarkably, 43 (68.3%) of the 63 early-life infant stool communities were dominated by *Bifidobacterium*, *Streptococcus*, or *Klebsiella* (**Fig. 6**d)—the same three genera highlighted in the South Africa infant stool metagenomes (**Fig. 5**d). We therefore tested the growth of representative strains across urea concentrations that spanned those encountered through human milk, infant formula, and higher-urea animal milk.

Bacterial responses were strongly dependent on strain and urea concentration (**Fig. 6**e–h). Some strains behaved as specialists, showing optimal growth within restricted concentration windows, whereas others benefited progressively as urea increased. Most notably, the opportunistic pathogen *Klebsiella pneumoniae* showed increasing growth with higher urea availability.

Together, these experiments show that feeding source can substantially alter infant urea exposure, with differences in urea availability capable of selectively favoring particular commensal or opportunistic members of the early-life ureabiome.

## Discussion

We identify urea as a conserved, dynamic component of mammalian milk with transport across the epithelium. Maternal stool and milk provide complementary reservoirs of candidate urea-metabolizing bacteria, and both milk and infant gut metagenomes encode urease and urea transport machinery. In overview, this constitutes a choreographed system of maternal-infant interaction facilitated by urea as a milk constituent. In contrast to breast milk, each infant formula contains a fixed level of urea, which does not vary dynamically; such differences can selectively affect important bacterial constituents of the early life intestinal microbiome^50,51^. Our findings connecting mammary urea handling with the metabolic ecology of the developing infant gut identify opportunities for improvements in infant nutrition.

Because urea is an end-product of protein catabolism, maternal protein intake and circulating urea are obvious candidate determinants of milk urea levels. Prior studies have tested dietary and systemic influences on urea during lactation, but controlled human and mouse interventions have yielded mixed results^52–54^. In our study, neither recent protein intake nor paired serum urea explained milk urea after adjustment for infant age and dietary covariates, and the same developmental trajectory occurred in mice maintained on a standardized diet. This instead points to controlled regulation within the mammary gland: AQP3—the only urea-permeable transporter^55^ robustly associated with the lactating secretory epithelium—provides a route compatible with blood-to-milk urea transfer, while loss of epithelial UT-B may restrict parallel equilibration or backflux. Such co-ordination, conserved across human populations and mice, is difficult to reconcile with passive metabolic spillover alone and instead points to mediated transport.

Once urea reaches milk, its biological consequences depend on which bacteria are positioned to use it. We found substantial restructuring of urea-metabolizing taxa in the infant gut, particularly *Bifidobacterium* and *Klebsiella*. Both encoded complete urea-utilization machinery, yet their responses diverged: a tested *B. longum* strain grew optimally at urea concentrations characteristic of early-lactation human milk, whereas *K. pneumoniae* benefited progressively as urea increased. Early-life events can further determine which organism occupies the urea-metabolizing niche: in our cohort, opportunistic pathogen *Klebsiella* was enriched after Cesarean delivery, consistent with previous research^56^. In a separate U.S. cohort of 412 infants^57^, *Bifidobacterium* deficiency was more common after C-section and was not rescued by breastfeeding; in these communities, urease metabolism was instead carried largely by members of Firmicutes and Proteobacteria. Thus, vaginal birth together with breastfeeding may provide a physiological urea niche for infant-adapted commensals. When this balance is disturbed, the same nitrogen resource may be exploited by less favorable taxa.

These contrasting scenarios raise a clinically important question: how can infant nutrition better preserve the microbial ecology supported by human milk? This may be particularly important for infants whose microbial communities are already altered by C-section or other early-life exposures, such as perinatal antibiotics^56,58^. Efforts to improve infant formula have increasingly incorporated individual bioactive components of human milk, particularly HMOs^59^. Yet HMOs alone do not determine which commensals establish: in a longitudinal study of breastfed infants^60^, maternal HMO composition did not explain *Bifidobacterium* species abundance. Interestingly, *B. longum* subsp. *infantis* was not prominent immediately after birth, but emerged from ∼10 weeks and reached its highest abundance between 10 and 25 weeks. Notably, this window overlaps with the rise toward the ∼20-week milk-urea maximum in our study, suggesting that microbial succession may reflect changing combinations of milk-derived substrates rather than any single nutrient. This dynamic exposure is not recapitulated by formula feeding, as the products we tested contained fixed and substantially higher urea concentrations than human milk.

We highlight three priorities for future research. First, non-protein nitrogen should be considered alongside other milk-derived microbial resources, including vitamins and trace metals^61,62^, in assessing infant nutrition needs. Second, we need to determine how this nutritional landscape changes as the microbiome matures. Our data indicate that *Blautia* begins expanding at ∼12–20 weeks and notably, urease-positive *Blautia* can salvage urea-derived carbon, rather than nitrogen, into acetate and, through microbial cross-feeding, butyrate—both short-chain fatty acids (SCFAs) with important roles in intestinal physiology^40,63^. Third, and most importantly, defining when the infant microbiome is most plastic may reveal critical windows in which nutrition can support healthy early-life microbial assembly, particularly given associations between perturbed early-life microbiome states and later allergy, asthma, and adiposity^64–66^.

## Online Methods

### Human cohort recruitment and sample collection

The study included three participant cohorts:

#### North America (USA-1) cohort 1

Between 2019 and 2022, 38 COVID-19– negative, full-term (≥ 37 weeks) mothers from New Jersey, New York, Pennsylvania, and Puerto Rico collected milk by hand expression or personal pump without prior breast sterilization, followed by freezing at home. Infants were 1 week to 12 months of age at the time of collection. Samples (*n* = 174) were retrieved within 48 h of collection, transported on dry ice, and stored at −80°C. Early-morning samples (∼06:00; *n* = 38) from all mothers, representing the first milk of the day, were used to assess infant-age trends and analyze the milk microbiome. A subset of 22 mothers provided a paired early-morning sample approximately one month after the initial collection (*n* = 44 across two timepoints) to quantify month-to-month stability. To assess diurnal variation, all 38 mothers provided samples at ∼6:00, ∼12:00, ∼18:00, and ∼24:00 on the initial collection day. Sample collection was conducted under Rutgers University IRB approvals Pro2018002781 and Pro2020002169, as well as University of Puerto Rico, Medical Sciences Campus IRB approval B2310120, as reported^27^.

#### New Jersey Kids Study (NJKS) cohort

This cohort comprised 63 full-term (≥ 37 weeks) mother–infant dyads enrolled in 2025 and 2026 in the NJKS. Maternal stool was self-collected during trimesters 1 (*n* = 14), 2 (*n* = 34), and 3 (*n* = 63). At the first postnatal collection, 4–20 weeks after delivery, maternal stool (*n* = 63), breast milk (*n* = 63), and infant stool (*n* = 63) were collected at home using study-provided kits, with optional maternal blood collection (*n* = 37). Participants washed their hands, wiped the breast and areola with a clean, damp cloth, and hand-expressed or pumped milk into a study-provided glass vial. Stool was collected using study-provided stool catchers and maternal blood was self-collected using a study-provided TAP Micro blood collection device (YourBio Health; Medford MA). Samples were refrigerated after collection and packed with frozen cold packs for retrieval from participants’ homes, with pickup within 24 h. Upon arrival, blood was centrifuged at 2,500 × g for 20 min at room temperature to separate serum, and all samples were stored at −80°C. 11 mothers provided a second breast milk sample at 9–12 months postpartum. Information on delivery mode (vaginal or cesarean), infant formula supplementation (ever or never exposed), and maternal diet was obtained from participant-completed questionnaires. Since urea is an end-product of protein metabolism, dietary analyses focused on animal (processed meat, beef/pork/lamb, fish, and dairy) and plant-protein foods (beans/lentils/chickpeas/tofu and peanut butter/nuts). During trimester 3, mothers reported how often they consumed these foods over the previous month. Responses were scored as reported^67,68^ (< 1 time per week = 0.5; once per week = 1; 2–4 times per week = 3; daily or nearly daily = 7; and ≥ 2 times per day = 14), and summed separately for animal- and plant-protein foods. At the first postnatal collection, mothers reported their food consumption during the previous 24 h; animal-protein intake was recorded as the number of different food products consumed and plant-protein intake was recorded as yes or no. The study was approved by the Rutgers University IRB Pro2023000278.

#### South Africa (SA) cohort

A cohort of 23 full-term (≥ 37 weeks) mothers (*l*_iving without HIV_ = 11; *l*_iving with HIV_ = 12) was randomly selected from the InFANT study^69^ at the Midwife Obstetric Unit in Khayelitsha, Cape Town, South Africa. All mothers delivered vaginally and their infants all were exclusively breastfed through 4 weeks postpartum. Mothers living with HIV were receiving antiretroviral therapy (and had undetectable viral load) and their infants received antiretroviral prophylaxis. At the 4-week hospital visit, breast milk was manually expressed without prior breast sterilization, and infant stool was collected. Samples were transferred into sterile tubes, transported on ice, and frozen at −20°C until processing. Ethical approval was granted by the University of Cape Town Human Research Ethics Committee and the South African Health Research Ethics Committee.

Across all human cohorts, milk samples could be collected at any point during a breast expression rather than from a complete expression, in line with prior work showing no significant within-feed variation in milk urea nitrogen concentration^70^.

### Mouse study design and sample collection

Conventional wild-type C57BL/6N mice were bred from a single in-house F0 grandparental pair to maintain microbiome homogeneity and housed under specific-pathogen-free (SPF) conditions at an AAALAC International-accredited facility. Mice were maintained on a 12:12-h light:dark cycle (lights on at 07:00) at 22 ± 1°C and 30– 70% relative humidity in individually ventilated polysulfone cages (Allentown LLC; Allentown NJ) with pelleted cellulose bedding (Biofresh; Patterson NY,) and provided with Enviropak enrichment (W.F. Fisher and Son; Branchburg NJ). Mice had *ad libitum* access to hyperchlorinated water via an automated watering system (Avidity Science; Waterford WI) and irradiated standard chow (5058 Irradiated Laboratory Rodent Diet; Purina, Richmond IN). Germ-free C57BL/6N mice were housed in flexible-film isolators at the Rutgers University Gnotobiotic Research Core Facility and provided autoclaved irradiated standard chow and autoclaved hyperchlorinated water.

After reaching sexual maturity, 8-week-old females underwent estrus synchronization by daily exposure to male cage bedding for one week, followed by co-housing with an age-matched male for 10 days. In total, the study comprised 88 females across reproductive stages, including 34 untreated conventional mice, 20 antibiotic-treated conventional mice, and 34 germ-free mice. During pregnancy, antibiotic-treated dams received neomycin (1 mg per 20 g body weight per day; *n* = 11) or vancomycin (0.4 mg per 20 g body weight per day; *n* = 9) by daily 100-µl oral gavage in sterile water from embryonic day (E)12 to E18.

Milk was collected from a subset of lactating dams, including conventional (*n* = 40 samples from 20 dams of which *n* = 6 untreated, *n* = 7 neomycin-treated, and *n* = 7 vancomycin-treated) and germ-free (*n* = 10 samples from 10 dams) mice, under isoflurane anesthesia after a 4-h-separation from pups. Chemical depilation of the areola was applied to facilitate access, and a subcutaneous 100 µL injection of oxytocin (20 IU/mL) used to induce milk letdown, with a second injection given when required. Milk was obtained by gentle mammary massage and collected into glass capillary tubes. Conventional dams contributed up to three milk samples between postnatal day (P)7 and P19 and were euthanized at P7 or P19, whereas germ-free dams were milked once between P10 and P15 followed by same-day euthanasia.

At sacrifice, the right abdominal mammary gland was divided into thirds and processed as follows: (1) snap-frozen for bulk RNA-seq; (2) incubated overnight in RNAlater (Thermo Fisher Scientific; Waltham MA) at 4°C, then the reagent removed and stored, until RT-qPCR was performed; and (3) fixed flat in histology cassettes in 10% neutral-buffered formalin for 48 h, washed 3 × 30 min in PBS, incubated for 30 min in 50% ethanol, and stored in 70% ethanol at room temperature. Blood was allowed to clot for 30 min and centrifuged at 2,000 × g for 20 min at room temperature to separate serum. All samples except histology specimens were stored at −80°C.

To assess early lactation, samples also were obtained from eight litters at P1. Because direct milking yielded insufficient volumes at this stage, one randomly selected pup per litter was dissected and stomach contents (“milk spots”) were collected as a proxy for early milk composition. All mouse work was conducted under IACUC protocols 201900013, 202100111, 201800253, and 201900032.

### Urea and total protein quantitation

Thawed human and murine milk samples were delipidated by brief vortex-mixing and centrifuging 250 µL aliquots (50 µL for murine milk) at 10,000 × g for 15 min at room temperature, as described^71^. The lipid layer was manually removed with a pipette tip, and the delipidation step repeated twice to ensure complete lipid removal. Aqueous supernatant (100 µL for human milk, 40 µL for murine milk) was used for downstream urea (mg/dL; Berthelot’s assay^72^) and total protein assays (µg/mL; BCA^73^ Protein Assay Kit; Thermo Fisher Scientific) using a SpectraMax® iD3 reader (Molecular Devices; San Jose CA). All assays were performed in technical duplicate, corrected for negative-control absorbance, averaged, and quantified using standard curves (R^2^ > 0.98). Quality-control thresholds required coefficients of variation (CoV) < 10% for high confidence and 10–30% for acceptable repeatability; samples with CoV > 30% were reprocessed. Infant formula, commercial cow and goat milk, plant-based beverages, and milk samples used in temperature-stability assays were delipidated as above before urea quantitation, whereas human and murine serum were quantified for urea directly. Samples were excluded because of failed delipidation (South Africa, *n* = 1 milk), or severe hemolysis (NJKS, *n* = 3 serum). Absolute urea concentrations (mg/dL) were used as the primary measure across all analyses. Within each human cohort and in comparisons with infant formula, findings were additionally confirmed using urea concentrations standardized to total milk protein (µg urea per mg total protein).

### Re-analysis of published scRNA-seq datasets from human breast tissue and milk

To investigate the potential mechanisms of local urea metabolism and transport in the human breast, publicly available single-cell RNA-sequencing (scRNA-seq) datasets were examined for the expression of urea-cycle and associated ornithine-metabolism genes (*NAGS*, *CPS1*, *OTC*, *ASS1*, *ASL*, *ARG1*, *ARG2*, and *OAT*), urea-permeable aquaglyceroporins (*AQP3*, *AQP7*, *AQP9*, and *AQP10*), and canonical urea transporters (*SLC14A1* and *SLC14A2*). Data were loaded, analyzed, and visualized using the *Scanpy* package^74^ (Version 1.9.6) in Python (Version 3.10.12).

For each original study^30,31^, raw counts were normalized to 10,000 counts per cell and natural-log transformed with a pseudocount of 1. Prior to analysis and data visualization, genes that were expressed in < 10% of cells across all cell types were excluded. This filtering removed the urea cycle genes *NAGS*, *CPS1*, *OTC*, and *ARG1*, as well as the candidate urea transport genes *AQP10*, *SLC14A1*, and *SLC14A2*.

First, human breast tissue was examined using the Integrated Human Breast Cell Atlas^30^, based on seven scRNA-seq studies^30,32–37^. For our primary analysis, the atlas was restricted to 24 healthy, adult female, non-lactating donors without germline breast cancer mutations, yielding 1,258,611 cells and 15,166 genes across 36 samples^30^. We then assessed the reproducibility of these findings across the seven individual scRNA-seq studies comprising the atlas^30,32–37^ (**Extended Data Table 1**).

Breast-milk-derived cells were re-analyzed using a longitudinal single-cell atlas spanning different stages of human lactation^31^. This integrated scRNA-seq dataset comprised 48,478 cells and 34,206 genes across 59 samples from 15 donors.

### Bulk RNA-seq of murine mammary tissue

For bulk RNA-sequencing (RNA-seq), total RNA was extracted from ∼40 mg snap-frozen mammary tissue using the miRNeasy Mini Kit (QIAGEN; Hilden, Germany), using the manufacturer’s recommendations with the following modifications for lipid-rich tissue. Samples were homogenized with eight 2.3-mm chrome-steel beads in 1 ml QIAzol at 2,000 rpm for 45 s and centrifuged at 15,000 × g for 5 min at 4°C. The upper lipid layer was removed and ∼800 µl of the underlying lysate transferred to a fresh tube. Chloroform (200 µl) was added, followed by vigorous mixing for 15 s, incubation for 3 min at room temperature, and centrifugation at 15,000 × g for 15 min at 4°C. The upper aqueous phase (∼400 µl) was transferred without disturbing the interphase and mixed with an equal volume of 70% ethanol. RNA purification then proceeded from step 6 of the manufacturer’s miRNeasy Mini animal-tissue protocol, with on-column DNase treatment (80 µl DNase I mixture for 15 min at room temperature; RNase-Free DNase Set; QIAGEN) before standard washing and elution. Following the final wash, columns were centrifuged for 1 min, and RNA was eluted in 35 µl RNase-free water after a 5-min incubation at room temperature. RNA concentration and purity were assessed by spectrophotometry (NanoDrop One; Thermo Fisher Scientific; Software Version 2.2.0.16). Extraction-negative controls were processed in parallel and yielded no measurable RNA by spectrophotometry; these controls were not submitted for sequencing.

Purified RNA underwent an additional DNase treatment before RNA integrity assessment (Agilent Technologies; Santa Clara CA). Across the 38 female mammary samples studied, RIN was 7.0 ± 0.3 (mean ± SEM) and DV200 was 86.9 ± 0.8%. Libraries were prepared using rRNA depletion with ERCC RNA Spike-In Mix (Thermo Fisher Scientific) according to the manufacturer’s protocol, and sequenced at Azenta (South Plainfield NJ) on an Illumina NovaSeq X (Version 1.3) using 2 × 150-bp paired-end reads, yielding 48.4 ± 1.5 million read pairs per sample, with 94.9 ± 0.1% of bases ≥ Q30.

### Bulk RNA-seq processing and analysis

Raw paired-end FASTQ files were assessed with *FastQC*^75^ (Version 0.12.0), and quality-control metrics were summarized using *MultiQC*^76^(Version 1.34). Transcript abundances were quantified against the *Mus musculus* GRCm39 reference genome (Release 108; mm39) using *kb-python*^77^ (Version 0.29.5) with *kallisto* expectation-maximization. Across the 38 samples, 87.2 ± 0.01% (mean ± SEM) of read pairs were pseudoaligned, yielding 42.3 ± 1.4 million transcript counts and 77,231 ± 769 detected transcripts/sample. One hundred bootstraps were generated/sample to estimate transcript abundance uncertainty and read-to-transcript ambiguity (RTA) overdispersion.

Analyses examined the same urea-cycle and transporter genes as in the human scRNA-seq datasets, with transporters expanded to include all murine aquaporins. Differential expression was analyzed at the gene level using *edgeR*^78^ (Version 4.10.1). Transcript counts were summarized using *tximport*^79^ (Version 1.40.0), yielding 24.4 ± 1.7 million gene counts and 19,354 ± 143 detected genes per sample. Samples were combined into a single *edgeR* DGEList with microbiome-status and reproductive-stage annotations, and genes with expression equivalent to ≥ 10 counts in a median-sized library in ≥ 3 samples were retained. Counts were further normalized using effective library sizes estimated by the trimmed mean of M-values (TMM) method^80^ and fitted with negative-binomial generalized log-linear models in *edgeR*. Samples were grouped by microbiome status (conventional or germ-free) and reproductive stage (non-pregnant/non-lactating [baseline], late pregnancy [E18], early lactation [P2–3], peak lactation [P7–12], or weaning [P13–19]), and specified contrasts tested using quasi-likelihood F-tests. *P*-values were adjusted for multiple comparisons using the Benjamini–Hochberg FDR procedure and considered significant if < 0.05.

To resolve bulk-tissue expression patterns within the mammary epithelium, we examined the CRUK Cambridge Institute mouse mammary epithelial scRNA-seq atlas^39^ using its online Shiny application, focusing on luminal epithelial populations across reproductive stages.

Reproductive stage-dependent genes conserved across microbiome status were defined as those differentially expressed relative to baseline (non-pregnant/non-lactating control) at one or more reproductive stages, with concordant expression patterns in conventional and germ-free mice. For clustering, fitted log_2_-transformed counts per million (log_2_CPM) were averaged by reproductive stage, baseline-centered, and standardized by standard deviation. Hierarchical agglomerative clustering was performed using Euclidean distance and Ward’s method^81^ implemented in *fastcluster*^82^ (Version 1.3.0), with the optimal number of clusters determined using the gap statistic^83^implemented in the *cluster*^84^ package (Version 2.1.8.3).

### Confirmatory RT-qPCR of murine mammary tissue

For confirmatory RT-qPCR, ∼20 mg RNAlater-preserved mammary tissue was blotted dry with Kimwipes (Kimberly-Clark Professional; Roswell GA) and homogenized as described above, using 700 µl QIAzol. RNA purification then proceeded according to the manufacturer’s miRNeasy Mini protocol (QIAGEN), with on-column DNase treatment, spectrophotometric assessment of RNA concentration and purity, and extraction-negative controls performed as described for bulk RNA-seq. cDNA was synthesized using the Verso cDNA Synthesis Kit (Thermo Fisher Scientific). RT-qPCR targets were selected based on relevance to urea transport and to validate three lactation-associated expression patterns observed by bulk RNA-seq: increased *Aqp3* expression, decreased *Slc14a1* (UT-B) expression, and persistently low or undetectable expression of *Slc14a2* (UT-A).

Primers (**Extended Data Table 2**) were designed in Primer-BLAST^85^ (*Mus musculus*, TaxID: 10090) to generate 100–220 bp intron-spanning amplicons covering all transcript variants, using standard criteria (18–22 bp, 50–60% GC, 3′ GC-clamp, low self-complementarity). Amplification of *Aqp3*, *Slc14a1*, *Slc14a2*_V1, and *Slc14a2*_V2 was confirmed using kidney tissue as a positive control. The reference genes *Gapdh* and *Rpl13a*^86^ amplified consistently across all 88 mammary samples. Primer efficiencies were required to be 70–120% with standard curve R^2^ > 0.95 in mammary tissue.

RT-qPCR was performed in technical duplicates using the QuantiNova SYBR Green Kit (QIAGEN) on the LightCycler® 480 II Instrument (Roche; Basel, Switzerland; Version 1.5.1.62 SP3). Cycling conditions were 95°C for 10 min; 40 cycles of 95°C for 45 s, 55°C for 60 s, and 72°C for 90 s; melt at 95°C for 5 s and 65°C for 60 s; final 40°C for 30 s; hold at 4°C. Each reaction contained cDNA corresponding to 200 ng of starting RNA. Reactions were repeated if mean crossing-point (CP) SD ≥ 1.0 or CoV ≥ 10%, or if amplification appeared in any no-template control (CP < 40).

Relative gene expression was normalized to the geometric mean of *Gapdh* and *Rpl13a*^87^, and calibrated to the mean expression in conventional non-pregnant/non-lactating female mice (baseline). Because *Slc14a2*_V1 and *Slc14a2*_V2 showed low or undetectable expression in conventional non-pregnant/non-lactating mammary tissue, the mean expression among samples with detectable amplification was used as the calibrator for these transcripts. Samples with CP ≥ 35 in ≥ 3 independent assays were considered below the detection threshold and assigned a relative-expression pseudovalue of 0.001 (ln = −6.91) for statistical analyses.

### Immunohistochemistry and immunofluorescence of murine mammary tissue

Fixation and storage of the histology specimens are described above. Based on the transcriptomic results and pilot AQP3 immunohistochemistry, localization analyses combined conventional and germ-free mammary tissues, and focused on early (P2–3), peak (P11–13), and late lactation (P19). AQP3 was examined by confocal immunofluorescence (*n* = 4 early, *n* = 4 peak, *n* = 3 late lactation; previously established reference tissue: rat kidney), AQP9 by immunohistochemistry (*n* = 2 per stage; positive control: rat liver^88^), UT-B and UT-A1 by confocal immunofluorescence (*n* = 4 early, *n* = 4 peak, *n* = 3 late lactation; positive control: rat kidney).

Unless specified, all procedures were carried out at room temperature. Formalin-fixed tissue was dehydrated for 2 h each in 70%, 96%, and 99% ethanol, cleared overnight in xylene, infiltrated with paraffin for 2 h, and embedded. Sections were cut at 2 µm, mounted on Superfrost Plus slides (Thermo Fisher Scientific), dried for 1 week at room temperature, heated at 60°C for 1 h, and deparaffinized overnight in xylene. Sections were rehydrated for 2 × 10 min in 99% ethanol, 2 × 10 min in 96% ethanol, and 10 min in 70% ethanol and rinsed in distilled water. Epitope retrieval was performed overnight at 60°C in TEG buffer (10 mM Tris, 0.5 mM EGTA; pH 9), followed by 30 min on ice and quenching of residual aldehydes with 50 mM NH₄Cl for 30 min.

Sections were blocked for 3 × 10 min in PBS containing 1% BSA, 0.2% gelatin, and 0.05% saponin and incubated overnight at 4°C with rabbit anti-AQP3 (1:800; AQP-003; Alomone Labs; Jerusalem, Israel), rabbit anti-AQP9 (1:50; 685AP^88^), rabbit anti-UT-B (1:100; 25962-1-AP; Proteintech; Rosemont IL), or rabbit anti-UT-A1 (1:100; abx448476; Abbexa; Cambridge, UK) diluted in PBS containing 0.1% BSA and 0.3% Triton X-100. Sections were washed for 3 × 10 in PBS containing 0.1% BSA, 0.2% gelatin, and 0.05% saponin.

For immunohistochemistry, endogenous peroxidase activity was blocked with 0.3% hydrogen peroxide in methanol, followed by incubation for 1 h at room temperature with horseradish peroxidase-conjugated goat anti-rabbit antibody (1:500; P0448; Agilent Technologies), DAB development (Agilent Technologies), Mayer’s hematoxylin counterstaining, dehydration, and mounting in DPX medium (Merck; Darmstadt, Germany).

For immunofluorescence, sections were incubated for 1 h at room temperature with Alexa Fluor 647- or 594-conjugated donkey anti-rabbit antibody (1:500; A32795 or A21207; Thermo Fisher) and Hoechst 33342 (2 µg/ml; Thermo Fisher Scientific), washed for 3 × 10 min in PBS containing 0.1% BSA, and mounted in Glycergel (Dako; Glostrup, Denmark) supplemented with 2.5% antifade reagent (Merck). Secondary-antibody-only controls, omitting the primary antibody, were included for all stainings.

For imaging, immunohistochemistry sections were acquired at 40× on a Nikon Ci-E automated microscope (Nikon Instruments; Melville NY) with a CFI Plan Apochromat Lambda 40× objective and Digital Sight 100 camera controlled by NIS-Elements D (Version 7.00.00, 64-bit).

Confocal images were acquired as 0.5-µm z-stacks on an Andor BC43 spinning-disk microscope (Andor Technology; Belfast, UK) using a 60× Plan Apo oil-immersion objective (NA: 1.4; working distance: 0.13 mm), a 4.1-megapixel, 16-bit sCMOS camera, and Fusion software (Version 2.7) with 1 × 1 binning, and 405-, 561-, and 638-nm excitation for Hoechst 33342, Alexa Fluor 594, and Alexa Fluor 647, respectively. Laser power, exposure time, detector gain, and offset were held constant within comparisons.

One section per mouse and 1–3 fields per section were imaged per mouse. Fields with tissue folds, sectioning artifacts, or poor focus were excluded. Apical, lateral, and basal epithelial domains were assigned from alveolar morphology and nuclei stains. All images were processed in Fiji/ImageJ^89^ (Version 1.54p).

### 16S SSU rRNA gene sequencing of stool and milk

For the USA-1 cohort, 16S library preparation was performed as part of a larger sequencing library and included early-morning milk samples (*n* = 36; 2/38 samples failed library prep), DNA extraction blanks (*n* = 9), and barcoding PCR no-template controls (*n* = 10). The NJKS 16S library included maternal stool collected across pregnancy and postpartum (*n* = 174), postnatal milk (*n* = 63), and infant stool (*n* = 63), with all samples matched according to the 63 mother–infant dyads. Negative controls comprised DNA extraction blanks (*n* = 15), PCR no-template controls (*n* = 22), unused at-home collection swabs (*n* = 4), and random participant home air samples (*n* = 5). Milk samples (*n* = 23) from the South Africa cohort were included in the NJKS 16S library. Mouse maternal stool samples collected longitudinally across gestation and lactation were processed using the same stool DNA extraction, V4 amplification, and sequencing workflow described below.

For milk samples, the delipidated supernatant and resuspended pellet were combined (∼100 µL), and used for DNA extraction with the DNeasy PowerSoil Pro Kit (QIAGEN) following the manufacturer’s protocol. Stool samples (∼20–30 mg) were processed with same kit’s 96-well format. DNA extraction negatives were included in each setup. DNA quality was assessed using NanoDrop One (1–200 ng/µL; 260/280 > 1.8). The V4 region of the 16S SSU rRNA gene was amplified using Earth Microbiome Project primers^90^ 515F^91^ and 806R^92^ with Maxima Hot Start PCR Master Mix (Thermo Fisher Scientific) under standard cycling conditions, including initial 94°C for 3 min; 35 cycles of 94°C 45 s, 50°C 1 min, 72°C 1.5 min; final 72°C for 10 min. No-template controls were processed at each molecular stage to monitor contamination. Paired-end, single-indexed PCR reactions were processed in triplicate and pooled, quantified with PicoGreen (Thermo Fisher Scientific) on a SpectraMax® iD3 reader, and purified with the QIAquick PCR Purification Kit (QIAGEN). Final libraries were quantified with the Qubit™ dsDNA HS Assay (Invitrogen; Waltham MA), diluted to 30 nM, and pooled equimolarly. The libraries were sequenced using 2 × 150-bp paired-end reads with a ∼10–15% PhiX spike-in. The USA-1 cohort library was sequenced on an Illumina MiSeq by Azenta and the combined library of NJKS and South Africa samples on an Illumina MiSeq i100 at the NYU Langone Genome Technology Center.

### 16S SSU rRNA gene sequencing bioinformatic processing

All computational pipelines were conducted in R (Version 4.3.3). The sequencing data were demultiplexed and paired-end FASTQ files were processed with *DADA2*^93^ (Version 1.32.0). Standard quality filtering parameters were applied, leading to the expected loss of ∼10% of sequencing reads. Error rates were calculated, ASVs were inferred using pseudo-pooling, and non-specific amplicons were removed, retaining merged, denoised reads of 250–256 bp, consistent with the expected V4 primer product length of ∼254 bp. Chimeric sequences were removed, and taxonomy assigned using the SILVA database^94,95^ (Version 138.1 for the USA-1 library; Version 138.2 for the combined NJKS–South Africa library) using the *DADA2* naïve Bayesian classifier; species-level assignments were added by exact sequence matching as available. A maximum likelihood phylogenetic tree was constructed using the GTR + G + I model with *DECIPHER*^96^ (Version 2.30.0) and *phangorn*^97^ (Version 2.12.1), as proposed^98^. Potential contaminants were identified and removed with *decontam*^99^ (Version 1.22.0), which flags amplicon sequence variants (ASVs) based on prevalence in control versus biological samples (threshold = 0.51 for the USA-1 library; 0.50 for the combined NJKS–South Africa library), removing 205 and 286 contaminant ASVs, respectively. ASVs that failed to classify at the phylum level, or that classified as chloroplast or mitochondria were removed prior to analysis^100^. Final mean ± SEM sequencing depth (reads/sample) and richness (observed ASVs/sample), respectively, were 24,818 ± 1,591 and 340 ± 55 for USA-1 milk; 57,753 ± 3,809 and 198 ± 7 for NJKS milk; 69,050 ± 1,582 and 454 ± 8 for NJKS maternal stool; 64,519 ± 2,479 and 217 ± 7 for NJKS infant stool; and 31,582 ± 3,506 and 173 ± 23 for South Africa milk.

### RT-qPCR of 16S SSU rRNA gene copy number

For USA-1 and NJKS samples with sufficient template DNA, total bacterial load was quantified by SYBR Green qPCR on a Roche LightCycler real-time PCR system (Roche; Version 1.5.1.62 SP3) using universal bacterial 785F (5′-GGMTTAGATACCCBGGTAGTC-3′) and 907R (5′-CCGTCAATTCMTTTGAGTTT-3′) primers and SYBR Green qPCR Master Mix (QIAGEN) according to the manufacturer’s protocol. DNA eluates were diluted 1:10 and assayed in duplicate using 1 µL template in 20-µL reactions. Absolute 16S rRNA gene copy number was determined from a 10-fold dilution series of a quantified 4,530-bp plasmid containing a single *Staphylococcus epidermidis* 16S rRNA gene fragment. Cycling comprised 95°C for 5 min followed by 40 cycles of 95°C for 10 s, 60°C for 20 s, and 72°C for 20 s, followed by melt-curve analysis from 65–97°C. Run quality was assessed from positive and negative controls, melt-curve specificity, and standard-curve performance (R² ≥ 0.98 and 90–110% amplification efficiency). Samples were re-assayed when CoV ≥ 10%. Bacterial load was reported as 16S rRNA gene copies per µL of DNA eluate after correction for dilution.

### 16S SSU rRNA gene sequencing data analyses

For NJKS analyses, singleton ASVs were removed before downstream analyses. β-diversity was calculated at the ASV level using Bray–Curtis dissimilarity on relative-abundance-transformed data with *phyloseq*^101^ (Version 1.46.0) and *vegan*^102^ (Version 2.6.6.1). Differences in community composition were tested by PERMANOVA with 9,999 permutations constrained within Mother_ID for repeated maternal stool analyses. Homogeneity of multivariate dispersion was assessed using PERMDISP.

11 candidate genera containing species or strains with reported or predicted urea-metabolizing capacity (*Bifidobacterium*, *Blautia*, *Collinsella*, *Enterobacter*, *Escherichia*–*Shigella*, *Klebsiella*, *Lactobacillus*, *Ligilactobacillus*, *Ruminococcus*, *Staphylococcus*, and *Streptococcus*^18,40–43^) were screened. Genera with mean relative abundance ≥ 0.05% and prevalence ≥ 10% in NJKS infant stool were retained, yielding nine genera for downstream analyses. Their relative abundances were compared across maternal third-trimester stool, postpartum stool, milk, and infant stool using log_10_-transformed abundances and mixed-effects models accounting for repeated samples within mother.

Maternal–infant ASV sharing was compared between candidate urea-metabolizing and other genus-assigned ASVs using presence–absence data, with sharing with each infant’s own mother compared with unrelated mothers using 10,000 randomized maternal assignments. Associations between birth mode and candidate genera in infant stool were tested after adjustment for infant age, formula exposure, and milk urea concentration.

For USA-1, taxa detected in > 50% of samples were retained prior to network construction, resulting in 82 species that represented 90.9% of the total relative abundance. Retained taxa were then agglomerated to the genus level, and co-abundance networks were constructed from early-morning milk samples (∼06:00; 36 mothers) separately for 1 week, 1–4 months, and 5–12 months postpartum. Correlations were inferred using *SparCC*^103^ implemented in *SpiecEasi*^104^ (Version 1.1.3), with 20 iterations and 100 bootstrap replicates. Significant (*P* < 0.05) associations with |cor| ≥ 0.2 were retained. Networks were visualized using *ggraph* (Version 2.2.1), and degree, betweenness centrality, closeness, and neighborhood connectivity were calculated using *igraph* (Version 2.0.3) and *tidygraph* (Version 1.3.1).

For mouse maternal stool microbiome data, community composition before and after antibiotic treatment was assessed at the ASV level using Jaccard dissimilarity calculated from relative-abundance-transformed data and ANOSIM with 9,999 permutations. Longitudinal changes in candidate urea-metabolizing genera were tested using linear mixed-effects models of arcsine-square-root-transformed relative abundance, with treatment, time, and their interaction as fixed effects and dam identity as a random intercept; Benjamini–Hochberg FDR correction was applied across genera.

### Shotgun metagenomic processing and analysis of human milk and infant stool

Shotgun metagenomic sequencing was performed on early-morning milk samples from the USA-1 cohort (*n* = 38). Libraries were prepared from genomic DNA using the NEXTFLEX Rapid XP V2 DNA-Seq Kit (Revvity; Waltham MA) on a Sciclone G3 NGSx iQ workstation, with PCR amplification and auto-normalization performed on an n6 iconPCR thermal cycler using the slope method. The library included DNA extraction negatives (*n* = 2), library-preparation negatives (*n* = 5), and a positive control (*n* = 1). Libraries were sequenced using 2 × 150-bp reads on an Illumina NovaSeq X with a 25B flow cell. Library preparation was performed at the Genomics and Microbiome Core Facility at Rush University (Chicago IL), and sequencing at the DNA Services Core, Roy J. Carver Biotechnology Center, University of Illinois Urbana-Champaign (Urbana IL). For the South Africa cohort, paired infant stool samples corresponding to the milk samples included in this study had previously undergone shotgun-sequencing as part of the InFANT study^69^. Of the 23 milk samples, metagenomic data were available for a subset of 19 infants, including 9 nursed by mothers living without HIV and 10 nursed by mothers living with HIV. All 19 infants were living without HIV.

All shotgun metagenomic analyses were performed using command-line workflows on a Linux server environment. Raw paired-end FASTQ files were processed with *KneadData*^105^ (Version 0.12.3) for adaptor trimming, quality filtering, and removal of host-derived reads. For USA-1 milk samples, quality-controlled reads were additionally taxonomically classified using *Kraken2*^106^ (Version 2.17.1) against the PlusPF database (k2_pluspf_20260226_16G), with species-level abundances re-estimated using *Bracken*^107^ (Version 3.1) with a read length of 150 bp. Potential contaminants were identified from taxonomic profiles using the prevalence method implemented in *decontam*^99^ (prevalence threshold = 0.1; Version 1.30), using the negative controls described above. Only reads passing KneadData filtering and not classified as human by Kraken2 were retained for subsequent processing.

Both USA-1 milk and South Africa infant stool metagenomes were subsequently processed with *SqueezeMeta*^108^ (Version 1.7.2) for assembly, annotation, and binning. Assemblies were generated using *MEGAHIT*^109^ (Version 1.2.9), and < 200 bp contigs removed and contig statistics generated with *PRINSEQ*^110^ (Version 0.20.4). Structural annotation included rRNA prediction with *Barrnap* (Version 0.9) and tRNA/tmRNA gene identification using *ARAGORN*^111^ (Version 1.2.38). Open reading frames (ORFs) were predicted with *Prodigal*^112^ (Version 2.6.3). Cleaned reads were mapped back to contigs using *Bowtie2*^113^ (Version 2.3.4.1) to estimate coverage.

Functional annotation of predicted proteins was conducted with *DIAMOND*^114^ (Version 2.0.15.153) against the *GenBank*^115^ nonredundant protein database (Release 2023.9), *eggNOG*^116^ (Version 5.0.2), and the Kyoto Encyclopedia of Genes and Genomes (*KEGG*^117^; Release 2011.6). KEGG Orthologs (KOs) were assigned from KEGG annotations, and Clusters of Orthologous Groups (COG) functional categories were derived from the *eggNOG* annotation layer. Additional hidden Markov model (HMM)-based homology searches were performed using *HMMER3*^118^ (Version 3.4) against the *Pfam*^119^ database (Version 35.0). Taxonomic classification of assembled 16S rRNA sequences was performed using the *RDP Classifier*^120^ (Version 2.10.2). USA-1 milk samples with < 10,000 KEGG-mapped reads were excluded from all downstream analyses (*n* = 6 removed; *n* = 32 retained). For KO-level analyses of USA-1 milk metagenomes, KOs were retained when coverage was > 1×, > 10 reads mapped to the KO, and prevalence was > 20% of retained samples (≥ 7/32).

Metagenome-assembled genomes (MAGs) were reconstructed using *MetaBAT2*^121^ (Version 2.12.1), and bin refinement was performed using *DAS Tool*^122^ (Version 1.1.1). MAG completeness and contamination were assessed with *CheckM2*^123^ (Version 1.0.2). Metabolic pathway inference for *KEGG* and *MetaCyc*^124^ (Release 2021.5) was performed using *MinPath*^125^ (Version 1.21) to obtain parsimonious pathway presence calls.

To characterize how urea metabolic functions are distributed across bacterial taxa, metagenomic reads mapping to urea-associated KOs were pooled within each cohort and assigned to bacterial genera based on contig taxonomy. For each urea-related KO, the relative contribution of each genus was calculated as the proportion of total reads mapping to that KO across the cohort. Urea-associated KOs, COG functional categories, and contributing bacterial genera were organized using unsupervised hierarchical clustering based on their relative contribution profiles. For bacterial taxa in which at least one urease structural gene (*ureA*, *ureB*, *ureC*) was detected by KEGG annotation, targeted follow-up analyses were performed to assess recovery of the complete *ureABC* structural module. ORFs annotated as urease-related were examined in contig context, and corresponding nucleotide sequences were extracted based on contig coordinates. These sequences were queried using BLASTX^126^ and BLASTN^126^ to confirm subunit identity and refine taxonomic assignment, particularly in cases of partial module recovery, split operons, or discrepancies between ORF-and contig-level taxonomy.

Final mean ± SEM sequencing depth (reads/sample), KEGG richness (observed KOs/sample), and species richness (observed species/sample), respectively, were 776,893 ± 308,175, 3,003 ± 221, and 307 ± 29 for USA-1 milk; and 123,605,962 ± 3,332,308, 6,963 ± 105, and 820 ± 14 for South Africa infant stool.

### Thermal stability of urea

To assess thermal stability of urea in different liquid environments, we tested aqueous urea solutions and delipidated human breast milk and whole cow milk (*n* = 8 per matrix; human milk samples were restricted to 1–4 months postpartum). Matched 20 µL aliquots from each sample were incubated for 30 min at five temperature conditions representing common storage or processing environments: 5°C (refrigeration), 20°C (room temperature), 37°C (physiological and warming before feeding), 61°C (pasteurization), and 100°C (boiling). Urea concentrations were measured in duplicate after thermal exposure, blank-corrected, and calculated from a standard curve (R² ≥ 0.99). Only measurements with CoV ≤ 10% were retained. Concentrations were corrected for volume loss during incubation where relevant, and log_2_-fold change was calculated relative to the corresponding 20°C condition, representing routine room-temperature handling.

### *In vitro* bacterial growth and urease activity in response to urea

Strains representing early-life-associated genera with reported or predicted ureolytic potential were selected for functional assays (**Extended Data Table 3**). The *Bifidobacterium* strains were isolated from infant stool collected in the ECAM (Early Childhood Antibiotics and Microbiome) Study^58^ by anaerobic culture on Bifidus Selective Medium (MilliporeSigma; Burlington MA) supplemented with BSM Supplement (MilliporeSigma), and identified by 16S rRNA gene Sanger sequencing followed by BLAST against the NCBI nucleotide database. Reference *Klebsiella pneumoniae* and *Streptococcus* strains were obtained from ATCC or the Cugini laboratory.

Growth assays included *Klebsiella pneumoniae*, *Bifidobacterium breve*, *B. bifidum*, *B. longum*, and *B. pseudocatenulatum*, and urease activity assays targeted *Streptococcus australis*, *S. cristatus*, *S. gordonii*, *S. parasanguinis*, *S. sanguinis*, and *S. sobrinus*. The latter was used for strains whose growth requirements necessitated additional nitrogen sources in minimal medium, precluding direct interpretation of urea-dependent growth. For both assays, urea concentrations were selected based on levels measured in this study, including a 0-mM control and concentrations representative of human milk (0.3, 0.6, and 0.9 mM), infant formula (1.2, 1.5/1.6, 1.8, and 3 mM), and cow milk (5 and 10 mM).

For growth assays, *K. pneumoniae* was recovered on LB agar (*n* = 4 biological replicates) and grown in LB for ∼16 h. *Bifidobacterium* strains were recovered on BSM agar (*n* = 6 biological replicates per strain) and grown anaerobically in mMRS with yeast extract for 24–48 h. Cells were then harvested, washed, and resuspended in the respective assay media. *K. pneumoniae* was assayed in nitrogen-free modified M9 medium^127^, whereas *Bifidobacterium* strains were assayed in mMRS without yeast extract and containing 0.03% L-cysteine^128^. Because L-cysteine could provide an additional nitrogen source, comparisons were made with the corresponding 0-mM urea control. Cultures were inoculated at a starting OD600 of ∼0.01 in 96-well plates at a final volume of 200 µL/well and incubated at 37°C in a BioTek LogPhase 600 reader. *K. pneumoniae* cultures were covered with gas-permeable membranes and shaken at 700 rpm, whereas *Bifidobacterium* cultures were sealed and incubated statically under anaerobic conditions. OD600 was measured every 10 min for 48 h and background-corrected against corresponding medium blanks. Growth was quantified as ΔOD600 by subtracting the initial OD600 at 0 h from the mean blank-corrected OD600 at 23–25 h for *K. pneumoniae* and the maximum OD600 at 4–6 h for *Bifidobacterium* strains.

Urease activity in *Streptococcus* strains was quantified using the phenol red assay modified from previous studies^129,130^. *Streptococcus* strains were recovered on BHI agar (n = 4 biological replicates per strain) and grown in BHI broth for ∼16 h. Cells were harvested, washed, and resuspended in M9-based minimal medium (pH 7.0) containing 6 g/L Na_2_HPO_4_, 3 g/L KH_2_PO_4_, and 0.5 g/L NaCl. Suspensions were adjusted to OD600 = 0.5 and inoculated into medium containing 0.1% (w/v) Casamino acids, 0.1% (w/v) phenol red, and the urea concentrations outlined above in 96-well plates at 200 µL/well. Cultures were incubated for 16 h at 37°C with shaking at 500 rpm. Absorbance at 430 and 560 nm was measured at 0 and 16 h using a SpectraMax® iD3 reader. Urease activity was calculated as the A560/A430 ratio at 16 h after subtraction of the corresponding 0-mM urea control ratio.

For each strain, the effects of urea concentration on bacterial growth or urease activity were assessed by one-way ANOVA, followed by Dunnett’s multiple-comparison test comparing each urea concentration with the corresponding 0-mM control.

### Statistical analysis and data visualization

Unless otherwise specified in the relevant Methods sections, statistical analyses were performed in R (Version 4.3.3). All hypothesis tests were two-sided. Distributional assumptions were evaluated using the Shapiro–Wilk test together with visual inspection of data distributions and model residuals, and homogeneity of variance was assessed using Levene’s test where applicable. Normally distributed continuous outcomes were compared using two-sample or paired t tests and analysis of variance (ANOVA), whereas Wilcoxon rank-sum, Wilcoxon signed-rank, Kruskal–Wallis, or Friedman tests were used for non-normally distributed data or matched observations as appropriate. When omnibus tests indicated significant effects, post-hoc comparisons were performed using Tukey’s honestly significant difference test following ANOVA, Dunn’s test following Kruskal–Wallis, or paired Wilcoxon signed-rank tests following Friedman tests. Benjamini–Hochberg false discovery rate correction was applied within prespecified families of related comparisons unless otherwise stated; Holm-adjustment was used for post-hoc comparisons following Friedman tests.

Associations between continuous variables were assessed using Pearson correlation when parametric assumptions were met and Spearman rank correlation otherwise. Positive continuous outcomes were additionally modeled using generalized linear models with a Gamma distribution and log-link where indicated, with exponentiated coefficients reported as ratios. HC3 heteroscedasticity-robust standard errors were used where indicated. Non-linear relationships between milk urea and infant age were evaluated using segmented linear regression with the breakpoint estimated from the fitted model.

For repeated-measures analyses, linear mixed-effects models were fitted using *lme4*^131^ (Version 1.1.35.5) and *lmerTest*^132^ (Version 3.1.3), with random intercepts for individual identity. Post hoc contrasts were estimated using *emmeans* (Version 1.11.2.8), with multiple-comparison adjustment as specified for each analysis. Model assumptions were assessed by inspection of residual distributions and Q–Q plots to confirm approximate normality and homoscedasticity; outcomes were ln- or log_10_-transformed where required to improve model fit.

Statistical significance was defined as *P* < 0.05. Figures were generated using *ggplot2*^133^ (Version 3.5.1) unless otherwise noted, and assembled and annotated using BioRender and Adobe Illustrator.

## Data Availability

All sequencing data and associated metadata generated in this study will be made openly available at the time of publication through widely used public repositories, including NCBI SRA and GEO, as appropriate. Previously published datasets analyzed here are publicly available as the Integrated Human Breast Cell Atlas through CZ CELLxGENE Discover (collection 48259aa8-f168-4bf5-b797-af8e88da6637), the longitudinal human breast milk single-cell RNA-seq atlas through the Broad Institute Single Cell Portal (SCP1671), the murine mammary gland single-cell RNA-seq dataset through the Mammary Gland Development portal and GEO (GSE106273), and the SA infant stool metagenomic dataset from the InFANT study through NCBI SRA (PRJNA1143669), with associated metadata.

## Code Availability

Custom R and Python code used for data processing, statistical analyses, and figure generation will be made openly available at the time of publication through public repositories, including GitHub and Zenodo. All third-party software, packages, and analytical methods are described in the Online Methods and cited where applicable.

## Acknowledgements

This paper is respectfully dedicated to the memory of Professor Tore Midvedt, whose enthusiasm for the conservation of urea in mammalian milk was the impetus for this study. We are grateful to Dr. Peter Agre for discussions on aquaporins; Dr. Andreas Riishede for pathological evaluation; Dr. Julie Halvorsen, Dr. Deborah Yao, and Dr. Piret Veerus for advice on early-life nutrition and infant formula; and Dr. Carla Cugini for sharing the *Streptococcus* isolates used in the bacterial assays. The research was supported by the New Jersey Health Foundation, the New Jersey Kids Study, the National Institutes of Health (U01AI122285), the C&D Fund, the Sergei S. Zlinkoff Fund for Medical Research and Education, and the Emch Fund for Microbiome, as well as by a CIFAR Fellowship to M.G.D.-B. and the Ferring Pharmaceuticals COVID-19 Investigational Grant in Reproductive Medicine and Maternal Health (RMMH). Additional support for D.D.N. and H.B.J. was provided by the National Institutes of Health (R01HD102239). A.A.A. and Q.N. were partially supported by the National Science Foundation (DMS-1763272) and the Simons Foundation (Grant 594598 to Q.N.).

## Author Contributions

L.V., E.R.L., and M.J.B. conceived the study. USA-1 human milk collection was performed by M.A.W., under the supervision of M.G.D.-B. Recruitment, sample collection and storage, and metadata development and management for the NJKS cohort were overseen by the New Jersey Kids Study team, including E.B., M.G.D.-B., and M.J.B., with laboratory coordination by M.T.-W. and Z.G. SA cohort sampling and metagenomic data generation were conducted by D.D.N. and H.B.J. Animal experiments were performed by L.V., J.E.S., E.R.L., D.S., and Y.S.Y., under the supervision of M.J.B. Urea and total-protein measurements were performed by D.S., R.C., and L.V. Re-analysis of human single-cell RNA-seq data was performed by A.A.A., under the supervision of Q.N. Bulk RNA-seq libraries were prepared by L.V., with processing and analyses performed by A.A.A. under the supervision of Q.N.; mammary RT-qPCR experiments were performed by D.S., E.R.L., and L.V. Mammary tissue immunohistochemistry and immunofluorescence were performed by V.S.D. and L.N.N. Human milk DNA extractions were performed by M.A.W., L.V., M.J.S., H.S., and M.T.-W., and stool DNA extractions by M.T.-W. 16S rRNA gene library preparation, bioinformatic processing, and analyses were performed by L.V.; total bacterial load quantitation was performed by B.C. and L.V. Co-abundance network and shotgun metagenomic analyses were performed by H.S. Urea temperature-stability experiments were performed by R.C., A.K., and L.V. Bifidobacterial strains were isolated from infant stool by X.S.Z. Bacterial growth and urease activity assays were performed by R.R.B. and V.C., under the supervision of M.M. Data visualization and statistical analyses were performed by L.V., A.A.A., and H.S. The manuscript was written by L.V. and M.J.B. All authors reviewed and approved the final manuscript.

