## Extended Data Tables and Figures for "Urea is a dynamic and functional component of mammalian breast milk"

### Extended Data Tables and Legends

**Extended Data Table 1. Summary and characteristics of the seven "Integrated Human Breast Cell Atlas"[30] scRNA-seq datasets included in the re-analysis of human breast tissue.** All included samples were from healthy, adult, non-lactating, cancer-mutation–free individuals. The dataset from Reed *et al.*[30] was used as the representative dataset due to its superior cell and gene coverage, while all seven studies were integrated to assess the consistency of findings across datasets.

| Dataset | Donor inclusion criteria | Donors | Cells | Genes |
|---|---|---|---|---|
| Pal *et al.* (2021)[32] | No germline breast cancer mutations | 17 | 98,719 | 15,165 |
| Gray *et al.* (2022)[33] | No germline breast cancer mutations | 3 | 16,337 | 15,164 |
| Murrow *et al*. (2022)[34] | Cancer- and mutation-free | 28 | 86,136 | 15,136 |
| Nee *et al*. (2022)[37] | No germline breast cancer mutations | 11 | 126,227 | 15,128 |
| Twigger *et al.* (2022)[35] | Non-lactating (tissue only) | 8 | 54,714 | 15,166 |
| Kumar *et al.* (2023)[36] | Mammoplasty tissue (mutation-free) | 76 | 518,684 | 15,166 |
| Reed *et al.* (2024)[30] | Mammoplasty tissue (mutation-free) | 24 | 357,794 | 15,166 |

**Extended Data Table 2. Primer sequences for RT-qPCR targeting selected murine aquaporin (*Aqp3*), urea transporter (*Slc14a1*, *Slc14a2*), and housekeeping genes (*Gapdh*, *Rpl13a*).** *Slc14a2* encodes six transcript variants with distinct exon structure; therefore, two primer sets were required: *Slc14a2*_V1 targets exons shared by five variants, while *Slc14a2*_V2 amplifies the unique sixth variant.

| **Gene (transcripts captured)** | **Forward primer (5' -> 3')** | **Reverse primer (5' -> 3')** |
|---|---|---|
| *Aqp3* (1/1) | CAGTTCATAGGCACAGCCGC | GGTGAAGAGGCGAGGTCCAA |
| *Slc14a1* (3/3) | CCTGTCAGCCACAGGACACT | GGAAAATGCCGCCTGTCCAC |
| *Slc14a2*_V1 (5/6) | GGCATGATCCTGGTGGCTCT | GAGTGCGTAGAACATGCCGC |
| *Slc14a2*_V2 (1/6) | AAAGAAGCCCAGGTGGCAGT | AGGGCAGTGGCTCTTTGGTT |
| *Gapdh* (8/8) | TGGCCTTCCGTGTTCCTAC | GAGTTGCTGTTGAAGTCGCA |
| *Rpl13a* (1/1) | GTACGCTGTGAAGGCATCAA | ATCCCATCCAACACCTTGAG |

**Extended Data Table 3. Bacterial strains used for functional urea assays and associated metadata.** Infant-derived *Bifidobacterium* strains were isolated from stool samples and are annotated by birth mode, predominant feeding mode, and infant age at collection. *Klebsiella pneumoniae* and *Streptococcus* strains were obtained from ATCC. * or ** indicates isolates originating from the same infant participant. NA – not available; ECAM – Early Childhood Antibiotics and Microbiome Study[58]; ATCC – American Type Culture Collection.

| **Species** | **Strain** | **ECAM or ATCC ID** | **Origin** | **Birth mode** | **Feeding mode** | **Age** | **Assay** |
|---|---|---|---|---|---|---|---|
| *Bifidobacterium bifidum* | #1* | ECAM C038.08 (D187) | Infant stool | Vaginal | Formula-dominant | 187 days (6.1 months) | Growth on urea |
| *Bifidobacterium bifidum* | #2 | ECAM C044.08 (D186) | Infant stool | Vaginal | Breast milk-dominant | 186 days (6.1 months) | Growth on urea |
| *Bifidobacterium breve* | #1** | ECAM C042.07 | Infant stool | Vaginal | Formula-dominant | 182 days (6.0 months) | Growth on urea |
| *Bifidobacterium breve* | #2** | ECAM C042.07 | Infant stool | Vaginal | Formula-dominant | 182 days (6.0 months) | Growth on urea |
| *Bifidobacterium longum* | #1* | ECAM C038.08 (D187) | Infant stool | Vaginal | Formula-dominant | 187 days (6.1 months) | Growth on urea |
| *Bifidobacterium longum* | #2 | ECAM C045.06 | Infant stool | Vaginal | Breast milk-dominant | 154 days (5.1 months) | Growth on urea |
| *Bifidobacterium pseudocatenulatum* | #1 | ECAM C041.08 | Infant stool | Vaginal | Breast milk-dominant | 186 days (6.1 months) | Growth on urea |
| *Bifidobacterium pseudocatenulatum* | #2 | ECAM C010.16 | Infant stool | Vaginal | Formula-dominant | 431 days (14.2 months) | Growth on urea |
| *Klebsiella pneumoniae* | NA | ATCC 13883 | NA | NA | NA | NA | Growth on urea |
| *Streptococcus australis* | NA | ATCC 700641 | NA | NA | NA | NA | Urease activity |
| *Streptococcus cristatus* | NA | ATCC 49999 | NA | NA | NA | NA | Urease activity |
| *Streptococcus gordonii* | NA | ATCC 10558 | NA | NA | NA | NA | Urease activity |
| *Streptococcus parasanguinis* | NA | ATCC 15909 | NA | NA | NA | NA | Urease activity |
| *Streptococcus sanguinis* | NA | ATCC 10556 | NA | NA | NA | NA | Urease activity |
| *Streptococcus sobrinus* | NA | ATCC 33478 | NA | NA | NA | NA | Urease activity |

### Extended Data Figures and Legends

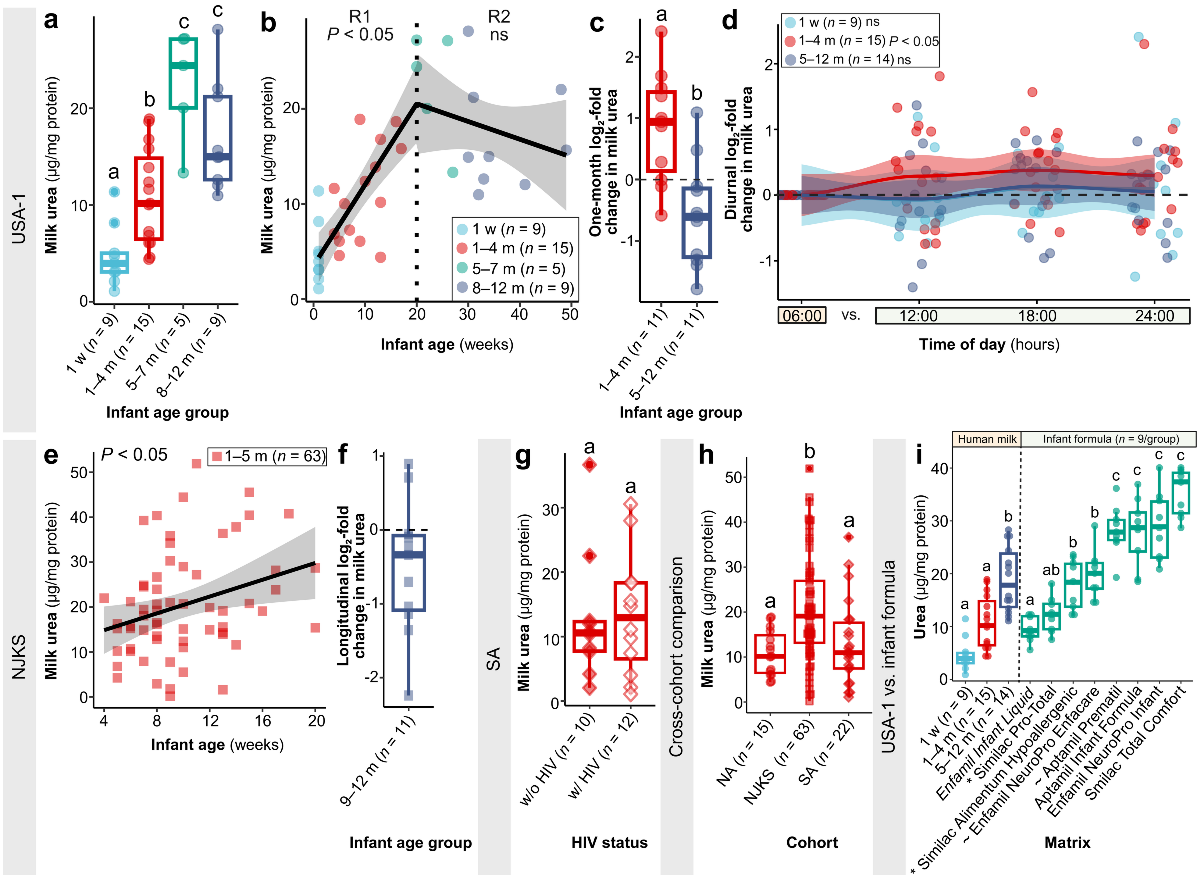


**Extended Data Figure 1. Protein-normalized milk urea levels recapitulate temporal, cross-cohort, and feeding-mode patterns.** Milk urea concentrations were normalized to total protein and are shown as µg urea per mg protein. Each point represents an individual milk sample. Groups assigned different letters differ significantly (P < 0.05 after multiple-comparison adjustment where applicable). (a) Protein-normalized milk urea levels across four infant age groups in 38 USA-1 mothers. (b) Segmented regression of normalized milk urea levels against infant age in USA-1. The dashed line marks the 20-week breakpoint and shading indicates the 95% confidence interval. (c) One-month $log_2$-fold change in normalized milk urea levels in paired USA-1 samples. (d) Within-day $log_2$-fold change in normalized milk urea levels relative to 06:00 in USA-1 mothers sampled at 6-h intervals. (e) Relationship between infant age and normalized milk urea levels in 63 NJKS mothers.

(f) $\log_2$-fold change in normalized milk urea levels from 1–5 to 9–12 months in 11 NJKS mothers with prolonged nursing. (g) Normalized milk urea levels in 22 South African mothers living without or with HIV. (h) Normalized milk urea levels across age-matched USA-1, NJKS, and SA cohorts. Unlike the comparison of absolute milk urea concentrations, significant pairwise differences emerged after total protein normalization, consistent with differences in mean total protein concentration among cohorts (USA-1: 3,358 ± 64 μg/mL (mean ± SEM); NJKS: 2,357 ± 86 μg/mL; SA: 4,050 ± 147 μg/mL). (i) Protein-normalized urea concentrations in human milk across lactation compared with nine commercial infant formula products per group. Annotation of formulas is identical with **Fig. 6**a. However, formula order differs slightly after total protein normalization because products vary in protein content. USA-1 – North America cohort 1; NJKS – New Jersey Kids Study; SA – South Africa cohort; w – week; m – months; w/o – without; w/ – with; ns – not significant.

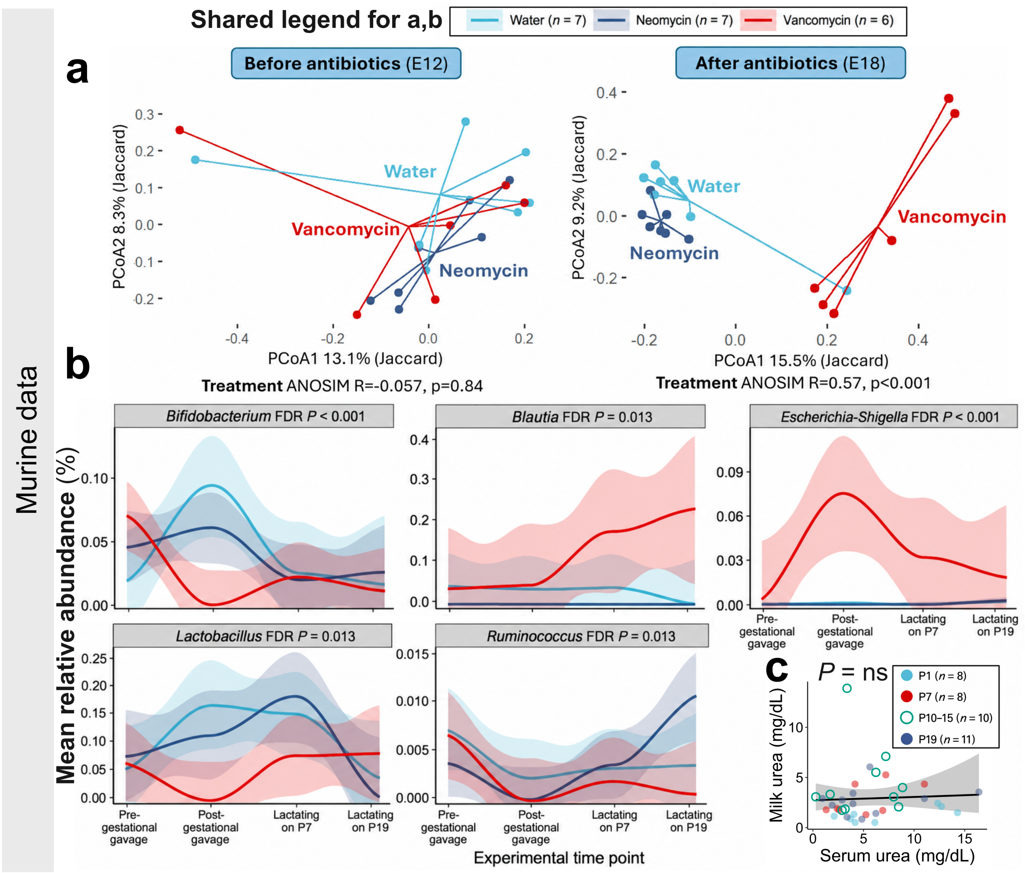


**Extended Data Figure 2. Maternal microbiome perturbation and circulating urea levels in mice.** (a,b) Jaccard-based PCoA of maternal stool microbiomes (a) before and (b) after antibiotic treatment. (b) Of the nine candidate ureabiome genera examined in the human analyses, five showed significant treatment-by-time interactions in maternal mouse stool after FDR-correction. Curves show LOESS fits with 95% confidence intervals; significance was assessed using linear mixed-effects models with dam as a random effect. (c) Paired serum and milk urea concentrations across murine lactation. Since serum was unavailable from conventional dams (filled circles) at peak lactation (P10–15), germ-free P10–15 dams (empty circles) were used as proxy since milk urea levels did not differ significantly between conventional and germ-free dams at peak lactation (**Fig. 1**i). The black line and shading show the unadjusted pooled Gamma regression and 95% confidence interval. E – embryonic day; P – postnatal day; PCoA – principal coordinate analysis; FDR – false discovery rate; ns – not significant.

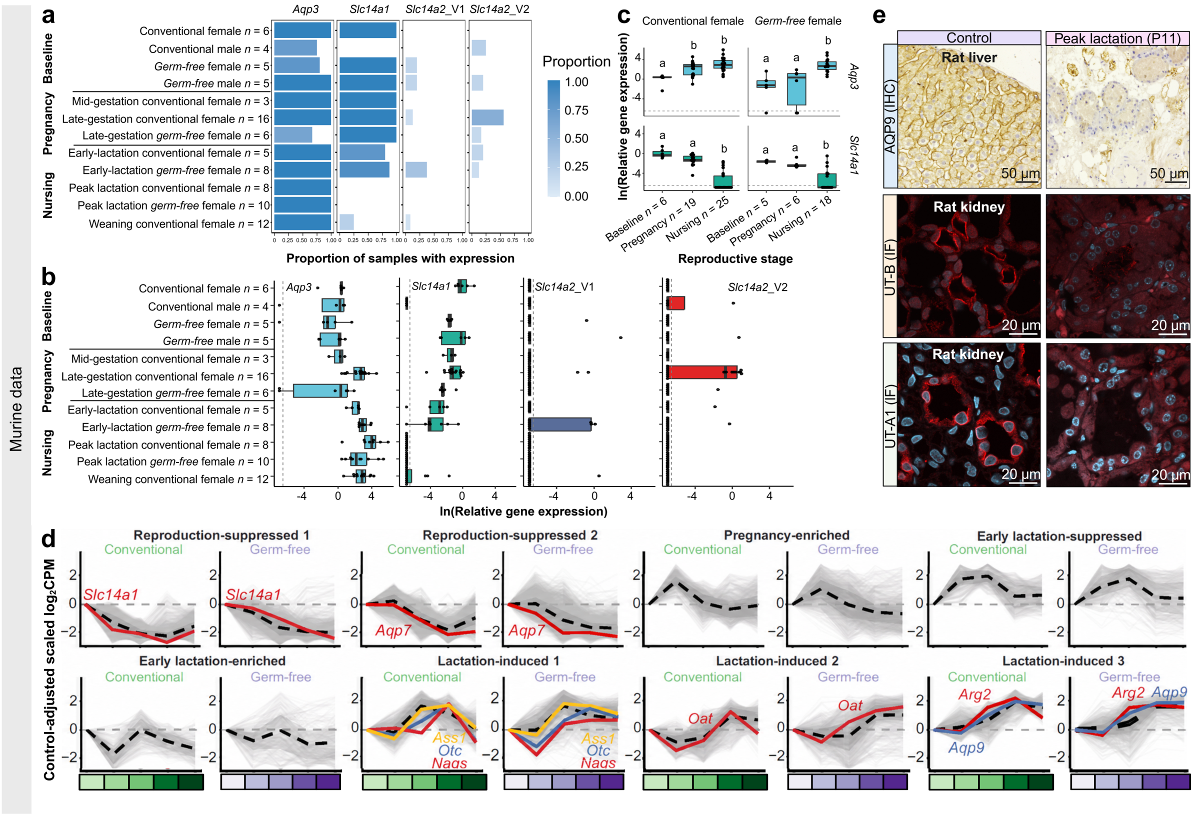


**Extended Data Figure 3. Reproductive stage-dependent regulation of mammary urea metabolism and transport in mice.** (a) Detection frequency of *Aqp3*, *Slc14a1*, and *Slc14a2* splice variants V1 and V2 across reproductive stages and microbiome states. (b) Corresponding ln-transformed relative expression among samples with detectable transcript. *Aqp3* and *Slc14a1* were normalized to non-pregnant, non-lactating conventional females; sparsely detected *Slc14a2* variants were normalized to the mean of positive samples. Dashed lines indicate detection thresholds. (c) *Aqp3* and *Slc14a1* expression after consolidation into baseline, pregnancy, and nursing stages; groups not sharing a letter differ significantly ($P$ < 0.05 after multiple-comparison adjustment where applicable). (d) Transcriptome-wide clustering identified eight additional reproductive stage-dependent expression programs. Urea cycle-associated genes, urea-permeable aquaporins, and canonical urea transporters are highlighted. The reproductive-stage colour gradient at the bottom of the panel denotes control, late pregnancy, early lactation, peak lactation, and late lactation, with greens for conventional and purples for germ-free mice. For conventional mice, $n$ = 3 at late pregnancy and peak lactation, and $n$ = 4 at all other stages; $n$ = 4 for all germ-free stages. (e) Representative immunohistochemistry (IHC) for AQP9 and immunofluorescence (IF) for UT-B and UT-A1 in respective positive control tissue and peak-lactation mammary tissue. AQP9: brown – immunoreactivity; blue-purple (hematoxylin) – nuclei. UT-B/UT-A1: red – immunoreactivity; blue (Hoechst) – nuclei.

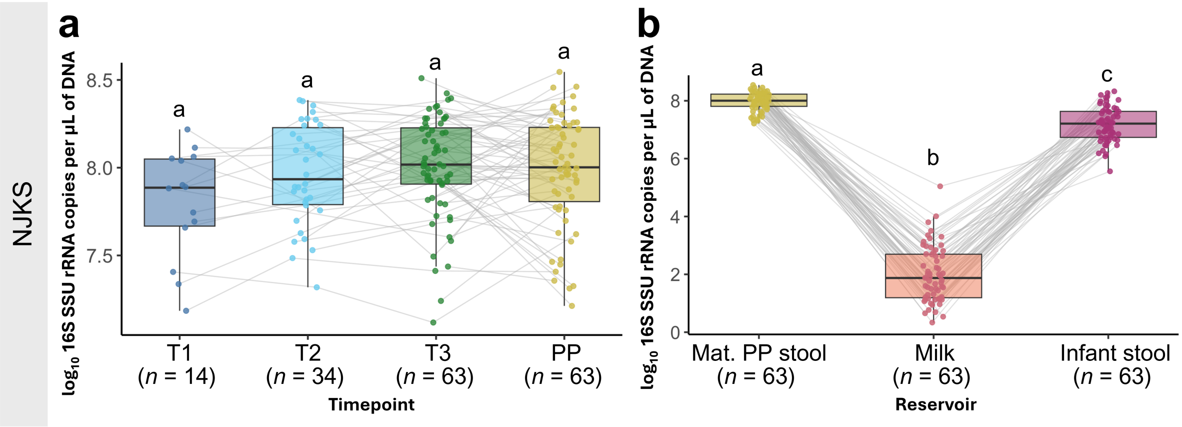


**Extended Data Figure 4. Total bacterial load across maternal and infant microbiomes in the NJKS cohort.** Groups not sharing a letter differ significantly ($P < 0.05$ after multiple-comparison adjustment where applicable). (a) Maternal stool bacterial load across pregnancy and postpartum did not differ by timepoint (linear mixed-effects model: $F_{3,\ 119.64} = 1.83$, $P = 0.15$). (b) Total bacterial load was significantly different among matched maternal stool, breast milk, and infant stool samples (Friedman test, $P < 0.001$). NJKS – New Jersey Kids Study; T1–T3 – first through third trimester; PP – postpartum.

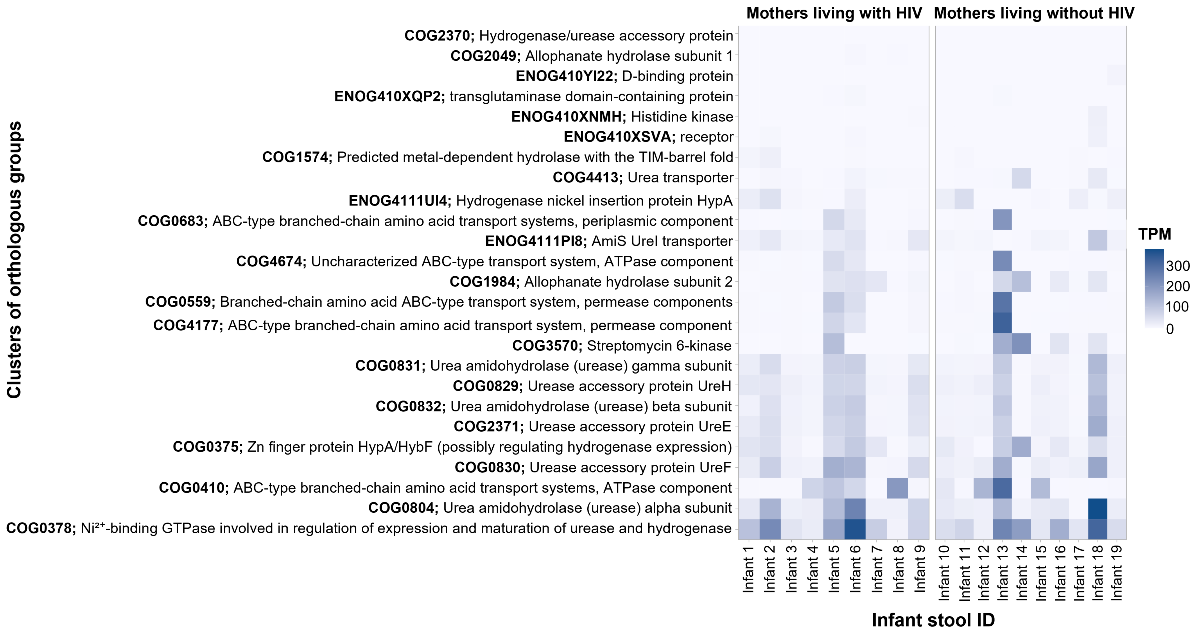


**Extended Data Figure 5. Urea- and nitrogen-associated functions in 19 SA infant stool metagenomes.** Heatmaps show the relative abundance of COGs related to urea metabolism and downstream nitrogen utilization in shotgun metagenomic data from stool samples of 4-week-old infants being nursed by mothers living with or without HIV. Color intensity reflects gene family abundance. SA – South Africa cohort; COG – Clusters of Orthologous Groups; TPM – transcripts per million.
